# Self-organization, quality control, and preclinical studies of human iPSC-derived retinal sheets for tissue-transplantation therapy

**DOI:** 10.1101/2022.02.18.480968

**Authors:** Kenji Watari, Suguru Yamasaki, Hung-Ya Tu, Masayuki Shikamura, Tatsuya Kamei, Hideki Adachi, Tomoaki Tochitani, Yasuyuki Kita, Aya Nakamura, Kazuki Ueyama, Keiichi Ono, Chikako Morinaga, Take Matsuyama, Junki Sho, Miyuki Nakamura, Masayo Fujiwara, Yoriko Hori, Anna Tanabe, Rina Hirai, Orie Terai, Osamu Ohno, Hidetaka Ohara, Tetsuya Hayama, Atsushi Ikeda, Daiki Nukaya, Keizo Matsushita, Masayo Takahashi, Akiyoshi Kishino, Toru Kimura, Shin Kawamata, Michiko Mandai, Atsushi Kuwahara

## Abstract

Three-dimensional retinal organoids (3D-retinas) are a promising graft source for transplantation therapy. We previously developed self-organizing culture for 3D-retina generation from human pluripotent stem cells (hPSCs). Here we present a quality control method and preclinical studies for tissue-sheet transplantation. Self-organizing hPSCs differentiated into both retinal and off-target tissues. Gene expression analyses identified the major off-target tissues as eye-related, cortex-like, and spinal cord-like tissues. For quality control, we developed a qPCR-based test in which each hPSC-derived neuroepithelium was dissected into two tissue-sheets: inner-central sheet for transplantation and outer-peripheral sheet for qPCR to ensure retinal tissue selection. During qPCR, tissue-sheets were stored for 3–4 days using a newly developed preservation method. In a rat tumorigenicity study, no transplant-related adverse events were observed. In retinal degeneration model rats, retinal transplants differentiated into mature photoreceptors and exhibited light responses in electrophysiology assays. These results demonstrate our rationale toward self-organizing retinal sheet transplantation therapy.

## Introduction

Pluripotent stem cells (PSCs) have the ability to self-organize three-dimensional (3D) neural tissue^1^. Self-organizing aggregates with 3D-tissue are called organoids, and represent promising graft sources for cell/tissue-transplantation therapy. Toward human retinal transplantation therapy, several retinal differentiation methods have been developed^2–22^. Among them, a self-organizing stem cell culture technology, SFEBq (serum-free floating culture of embryoid body-like aggregates with quick aggregation), is useful for 3D-retina generation^5–8, 15, 17^. We previously modified the SFEBq method by timed bone morphogenetic protein (BMP) treatment to generate 3D-retinas from feeder-free human embryonic stem cells (ESCs) and induced pluripotent stem cells (iPSCs)^7, 8^. The 3D-retinas in this culture system contain neural retinal (retinal) progenitors that expand to give rise to photoreceptors and other retinal neurons and form a multilayered stratified retinal tissue in a stage-dependent manner, recapitulating the development *in vivo*.

Retinitis pigmentosa is a group of hereditary diseases characterized by progressive loss of rod photoreceptors followed by cone photoreceptors, and is the major cause of blindness in developed countries. Transplantation of photoreceptors and/or retinal progenitors is a promising therapeutic option^23–26^. Indeed, transplantation of retinal tissues and cells from primary embryonic tissues had the potential to improve visual function^23, 27–29^, although the contributions of tissue/cell engraftment and donor-host material transfer remain to be studied^30^. To supply adequate amounts of retinal tissues and cells for cell therapy, 3D-retinas generated from PSCs represent a promising graft source. Photoreceptor precursors purified from PSC-derived retinas can survive, make contact with the host retina, and improve visual function^10, 17, 18, 26, 31, 32^.

Mouse and human retinal tissue sheets (retinal sheets hereafter) dissected from 3D-retinas became engrafted into end-stage retinal degeneration model mice and rats and exhibited functional maturation that resulted in restoration of light responses^33–37^.

Furthermore, hPSC-derived 3D-retinas had low immunogenicity and immunosuppressive properties^38^. Transplantation of human PSC-retinal sheets in primate models of retinal degeneration showed long-term engraftment for 2 years^36, 39^.

These preclinical transplantation studies indicate that allogeneic PSC-derived retinal sheets have the potential to improve visual function.

Toward clinical applications on PSC-derived retinal sheets, the establishment of a quality control (QC) strategy for 3D-retinas (intermediate products) and dissected retinal sheets (final products) has remained a major challenge. Self-organizing culture has been improved to reduce the variation in organoids at both the intra-batch and inter-batch levels. Eiraku and colleagues developed the SFEBq method to regulate the size and quality of PSC-derived aggregates^5, 40^. We developed a modified culture system to reduce the variety of organoid quality, such as the organoid morphologies and proportions of neural retina (retina) and retinal pigment epithelium (RPE)^7, 8^, although individual organoids still exhibit diverse 3D morphologies and multiple off-target tissues. Some off-target tissues were identified as RPE tissue and cortex-like tissue^7, 12^.

Our previous transplantation studies used PSC lines harboring fluorescent protein knock-in reporter lines for Rx (also called Rax) and Crx to label retinal cells suitable for transplantation^6, 35, 39^. However, xenogeneic fluorescent protein knock-in reporter lines are not ideal for clinical applications. Here we establish a new QC strategy to select retinal tissue in the organoids. We then performed preclinical safety and efficacy studies to demonstrate the rationale for allogeneic retinal sheet transplantation therapy.

## Results

### Self-organizing culture of human iPSCs for 3D-retina and dissected retinal sheet generation

Toward retinal tissue-transplantation therapy, we developed a robust retinal differentiation method consisting of five processes based on previous studies^5–8, 33, 39^: 1) maintenance culture with feeder-free hPSC culture and preconditioning methods, 2) retinal differentiation with SFEBq and BMP methods, 3) induction-reversal culture, 4) maturation culture, and 5) dissection (Fig. 1a).

**Fig. 1.**
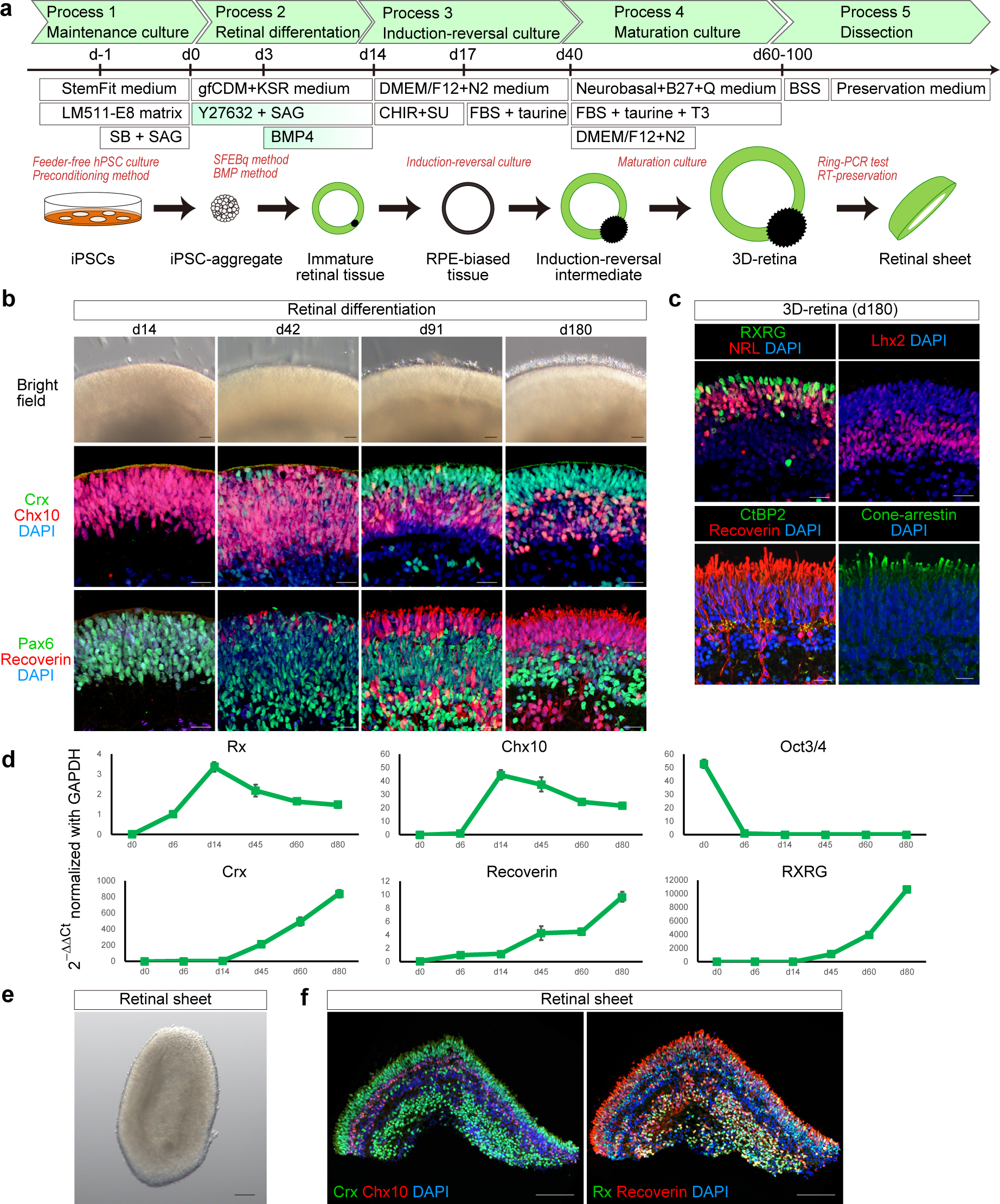
Self-organizing culture of human iPSCs to generate the 3D-retina and dissected retinal sheet. **a** Scheme of the self-organizing culture. **b** Bright-field view of iPSC-S17-derived cell aggregate containing retinal tissue on days 14, 42, 91, and 180 (upper). Scale bar in bright-field view: 100 µm. Immunostaining of iPSC-S17-derived retinal tissue on days 14, 42, 91, and 180 (middle and lower). Crx (green) and Chx10 (red) in middle panels. Pax6 (green) and Recoverin (red) in lower panels. Blue: nuclear staining with DAPI. Scale bar in immunostaining: 20 µm. **c** Immunostaining of iPSC-S17-derived retinal tissue on day 180. RXRG (green) and NRL (red) in upper-left panel. Lhx2 (red) in upper-right panel. CtBP2 (green) and Recoverin (red) in lower-left panel. Cone-arrestin (green) in lower-right panel. Blue: nuclear staining with DAPI. Scale bar: 20 µm. **d** Gene expression analysis in iPSC-S17-derived cell aggregates containing retinal tissue on days 0, 6, 14, 45, 60, and 80. In each replicate, RNA was extracted from 48 aggregates. mRNA levels were determined by qPCR analysis. Relative mRNA expression was determined by the delta-delta Ct method with *GAPDH* as an endogenous control. Data are presented as mean ± SE (*n* = 4 per time point). **e** Bright-field view of iPSC-S17-derived retinal sheet. Scale bar: 100 µm. **f** Immunostaining of iPSC-S17-derived retinal sheet on day 87. Crx (green) and Chx10 (red) in left panel. Rx (green) and Recoverin (red) in right panel. Blue: nuclear staining with DAPI. Scale bar: 100 µm.

In Process 1, human iPSC cell lines were maintained on LM511-E8 matrix in StemFit medium^41, 42^. Four iPSC cell lines, iPSC-S17, iPSC-LPF11, iPSC-1231A3, and iPSC-QHJI01s04 (iPSC-Q), were stably expanded in this feeder-free maintenance culture, and their doubling time was approximately 12–18 h. iPSCs were preconditioned with SB431542 (SB; inhibitor of TGF-beta/Nodal/Activin signaling) plus smoothened agonist (SAG; Shh signaling agonist) for 18–30 h prior to differentiation (SB+SAG hereafter)^8^. In Process 2, 3D-retinal differentiation culture was carried out using the SFEBq-BMP method. iPSCs were dissociated into single cells, plated in V-bottom 96-well plates, and cultured in differentiation medium (growth factor-free CDM [gfCDM] + Knockout Serum Replacement [KSR]) in the presence of Y-27632 and SAG^8^. The resulting iPSC-aggregates were treated with BMP4 on day 3 and cultured in gfCDM+KSR medium^7^. On day 14, the iPSC-aggregates were fixed and immunostained for retinal progenitor markers Pax6 and Chx10 (also called Vsx2). The iPSC-aggregates were found to contain a neuroepithelium on their surface, and the majority of the cells in the neuroepithelium were positive for Chx10 and Pax6, indicating that iPSCs self-formed immature retinal tissue (Fig. 1b). In Process 3, the immature retinal tissue was cultured using an induction-reversal culture method to precisely regulate the proportions of neural retina (retina) and RPE^7^. The immature retinal tissue was cultured with CHIR99021 (CHIR; Wnt agonist) and SU5402 (SU; FGFR inhibitor) for 3 days to bias the cells toward the RPE fate and then further cultured in retina maturation medium containing serum and taurine for 23 days to obtain an induction-reversal intermediate.

In Process 4, the induction-reversal intermediate was further cultured in retina maturation medium containing serum, taurine, and T3 for 20–60 days to obtain the 3D-retina. At days 60–100, the 3D-retina was fixed and immunostained for Chx10, Pax6, Crx (photoreceptor precursor marker), and Recoverin (photoreceptor marker).

The 3D-retina was found to contain a multilayered retinal tissue, comprising a photoreceptor precursor layer with Crx^+^ and Recoverin^+^ cells on the outer surface, a retinal progenitor layer with Chx10^+^/Pax6^+^ cells in the middle, and a neuronal layer with Chx10^−^/Pax6^++^ cells and Crx^+^ and Recoverin^+^ cells on the inner side (Fig. 1b and Supplementary Fig. 1a, b, c).

We further cultured 3D-retinas in long-term maturation medium containing all-trans retinoic acid to examine the expression of retinal maturation markers.

3D-retinas on day 180 were confirmed to contain a multilayered retinal tissue with photoreceptor and retinal progenitor layers (Fig. 1b, c). NRL^+^, RXRG^+^, and Cone-arrestin^+^ cells were found in the outer photoreceptor layer of the 3D-retina, indicating differentiation into rod precursors and cone precursors (Fig. 1c). Lhx2^+^ neuronal layer was found on the inner side of the 3D-retina (Fig. 1c). Some Recoverin^+^ cells expressed CtBP2 (also called Ribeye), suggesting that iPSC-derived photoreceptors had the ability to express photoreceptor synaptic proteins (Fig. 1c and Supplementary Fig. 1d).

To analyze gene expression, we performed qPCR analyses of iPSC-aggregates on days 0, 6, 14, 45, 60, and 80 (Fig. 1d and Supplementary Fig. 1e). The mRNA levels of retinal progenitor markers (*Rx*, *Chx10*), photoreceptor markers (*Crx*, *Recoverin*) and a cone photoreceptor precursor marker (*RXRG*) were increased at day 6–14, on day 45, and on day 60, respectively. The mRNA level of PSC marker *Oct3/4* was decreased on day 6. These data are consistent with previous 3D-retinal differentiation culture studies^5–8^.

In Process 5, 3D-retinas at days 60–100 were dissected into retinal sheets (final products) (Fig. 1e)^33, 39^. We checked the morphology of the retinal sheets by immunohistochemical (IHC) analysis and observed a multilayered continuous retinal tissue with a photoreceptor precursor layer containing Crx^+^ and Recoverin^+^ cells in the outer surface and a retinal progenitor cell layer with Chx10^+^ and Rx^+^ cells on the inner side (Fig. 1f). Using these five processes, human allogeneic iPSC-derived retinal sheets were reproducibly generated.

### Characterization of major off-target tissues in the retinal differentiation culture

The self-organizing culture mimicked embryonic development to induce both retinal tissue and off-target tissues. We sought to identify the major off-target tissues in this culture based on the morphology of the aggregates. iPSC-LPF11 cells differentiated toward the retinal fate by self-organizing culture were subjected to bright-field microscopy analysis. We found that iPSC-LPF11 cells self-organized into retinal tissue and three off-target tissues that had different morphologies from the retinal tissue (off-target tissue-1, -2, and -3) (Fig. 2a). The typical morphology of the retinal tissue was the multilayered continuous neuroepithelium structure with a bright outer layer and a brown inner layer (Figs. 1b and 2a). The typical morphologies of the off-target tissues were pigmented and/or non-pigmented thin-wrinkled epithelium (off-target tissue-1), sticky cell aggregates with a rough outer surface (off-target tissue-2), and dark-colored aggregates with a clogged inner part (off-target tissue-3) (Fig. 2a). We differentiated three independent culture batches from two cell lines, iPSC-LPF11 and iPSC-S17, and examined the proportions of aggregates containing retinal tissue and off-target tissues (Fig. 2b, *n* = 59–95 aggregates per each batch). For all batches, most aggregates contained retinal tissue and the proportion of aggregates containing off-target tissue-1 tended to be higher than the proportion of aggregates containing off-target tissue-2 or off-target tissue-3.

**Fig. 2.**
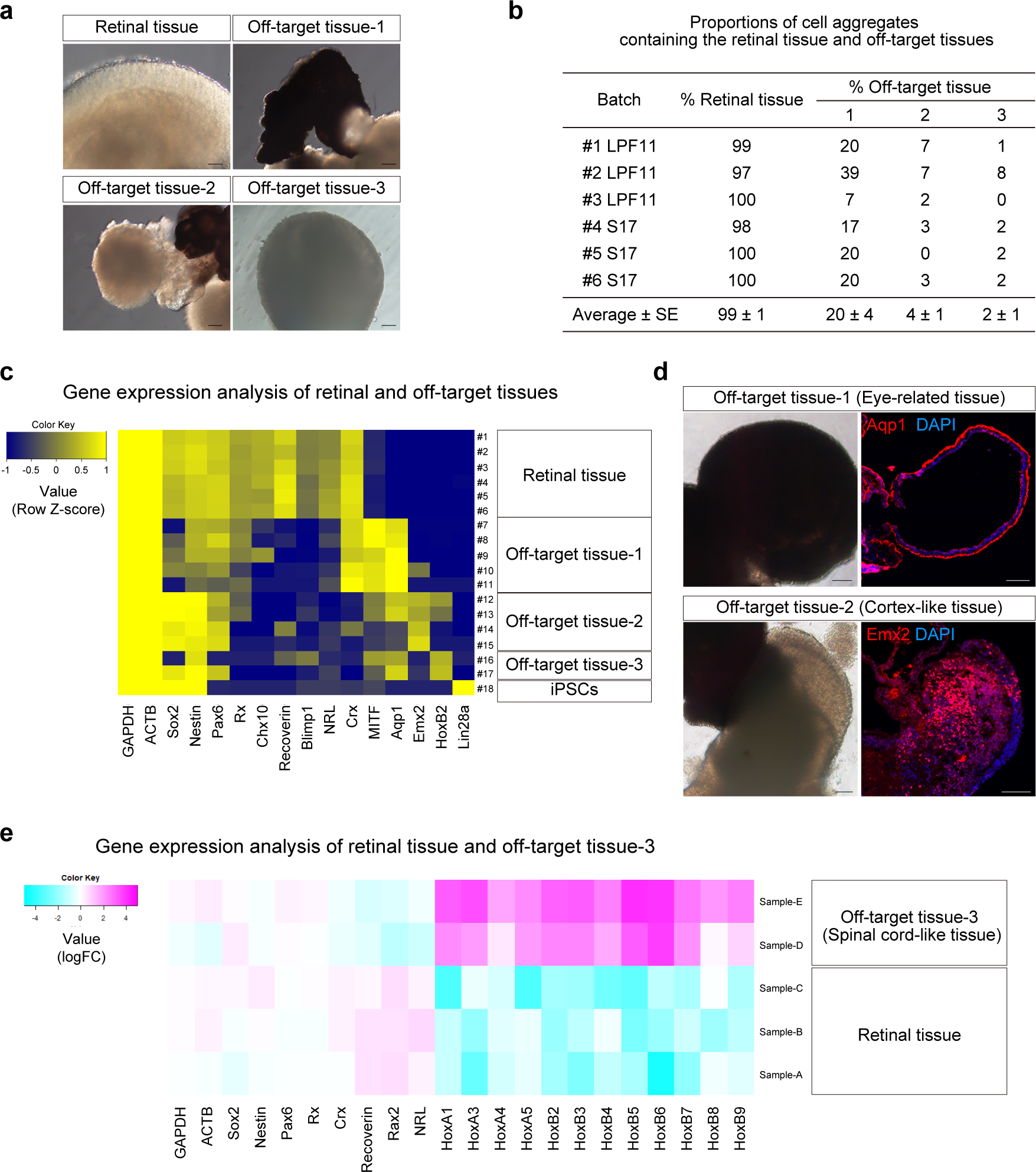
Characterization of major off-target tissues in the retinal differentiation culture. **a** Bright-field view of retinal tissue and off-target tissues in iPSC-LPF11-derived cell aggregates. Scale bar: 100 µm. **b** Proportions of cell aggregates containing retinal tissue and off-target tissues. Three independent batches derived from iPSC-LPF11 cells and three independent batches derived from iPSC-S17 cells were analyzed (*n* = 59–95 aggregates per each batch). Data are presented as mean ± SE. **c** Heat map for gene expressions in retinal tissue and off-target tissues derived from iPSC-LPF11 cells. As a control, gene expressions in undifferentiated iPSC-LPF11 cells are also shown. Gene expressions were measured by qPCR analyses. Row Z-scores were calculated for each tissue and plotted as a heat map. **d** Representative bright-field images and IHC images for off-target tissue-1 and -2 derived from iPSC-LPF11 cells. The bright-field images and IHC images were captured at the same positions. Immunostaining for Aqp1 (upper right; red) and Emx2 (lower right; red) is shown. Blue: nuclear staining with DAPI. Scale bar: 100 µm. **e** Heat map for gene expressions in retinal tissue and off-target tissue-3 derived from iPSC-1231A3 cells. mRNA levels were determined by microarray analysis. LogFC values were calculated for each gene and plotted as a heat map. FC, fold change. Note that off-target tissue-3 expressed Hox genes compared with retinal tissue.

To identify the cell lineages, iPSC-derived off-target tissue-1, off-target tissue-2, and off-target tissue-3 were manually dissected, possibly with small amounts of surrounding tissues, and the mRNA levels of key genes associated with fate choice, differentiation, and/or patterning were determined by qPCR analysis. Based on the Ct values for each gene, we calculated Z-scores for each tissue and plotted as a heat map to analyze the gene expression patterns in the tissues (Fig. 2c). We found that off-target tissue-1 expressed *Aqp1* and *Mitf*, indicating that it was eye-related tissue with RPE and ciliary body^43, 44^. Off-target tissue-2 expressed *Emx2*, indicating that it was embryonic cortex-like tissue^45^. Off-target tissue-3 showed high expression of homeobox gene *HoxB2*^46, 47^. Undifferentiated hPSC marker *Lin28a* was not detected in the three off-target tissues or retinal tissue^48^.

We performed IHC analyses and confirmed that off-target tissue-1 was positive for RPE and ciliary body marker Aqp1, while off-target tissue-2 was positive for embryonic cortical (dorsal telencephalic) marker Emx2 (Fig. 2d and Supplementary Fig. 2a). We investigated the gene expression in off-target tissue-3 by microarray analysis and found that it expressed multiple Hox genes including *HoxB2*, indicating that it was posterior-neural spinal cord-like tissue (Fig. 2e). We further confirmed that the gene expression patterns in the off-target tissues differed from those in the retinal tissues (Supplementary Fig. 2b).

Collectively, using self-organizing culture method, hPSCs robustly differentiated to form a multilayered retinal tissue and the major off-target tissues were Aqp1^+^ eye-related tissue (off-target tissue-1), Emx2^+^ cortex-like tissue (off-target tissue-2), and HoxB2^+^ spinal cord-like tissue (off-target tissue-3).

### QC method for retinal sheets by testing gene expression in the surrounding outer tissue-sheet

The self-organized retinal tissue had a unique multilayered continuous neuroepithelium structure (Figs. 1 and 2). Therefore, we can identify the retinal tissue by carefully observing the aggregates under a bright-field microscope during the dissection (Process 5). However, PSCs in this culture differentiated into not only retinal tissue but also off-target tissues (Fig. 2), providing a potential risk that off-target tissues will be dissected to generate the final product. Toward clinical applications, we aimed to develop a fail-safe QC method to ensure the selection of transplantable retinal sheets and avoid including off-target tissues by mistake. Although the qPCR-based QC test for all retinal sheets is a sensitive way to detect off-target tissues, execution of this test leads to the destruction of all transplantable retinal sheets. To avoid this destruction, we decided to dissect the continuous neuroepithelium into two tissue-sheets: the inner-central sheet for transplantation (named ‘cap’) and the outer-peripheral sheet for QC test (named ‘ring’) (Fig. 3a). To estimate the gene expression of each inner-central sheet (cap), we carry out qPCR analysis of each corresponding outer-peripheral sheet (ring) (Fig. 3a; ring-PCR test hereafter).

**Fig. 3.**
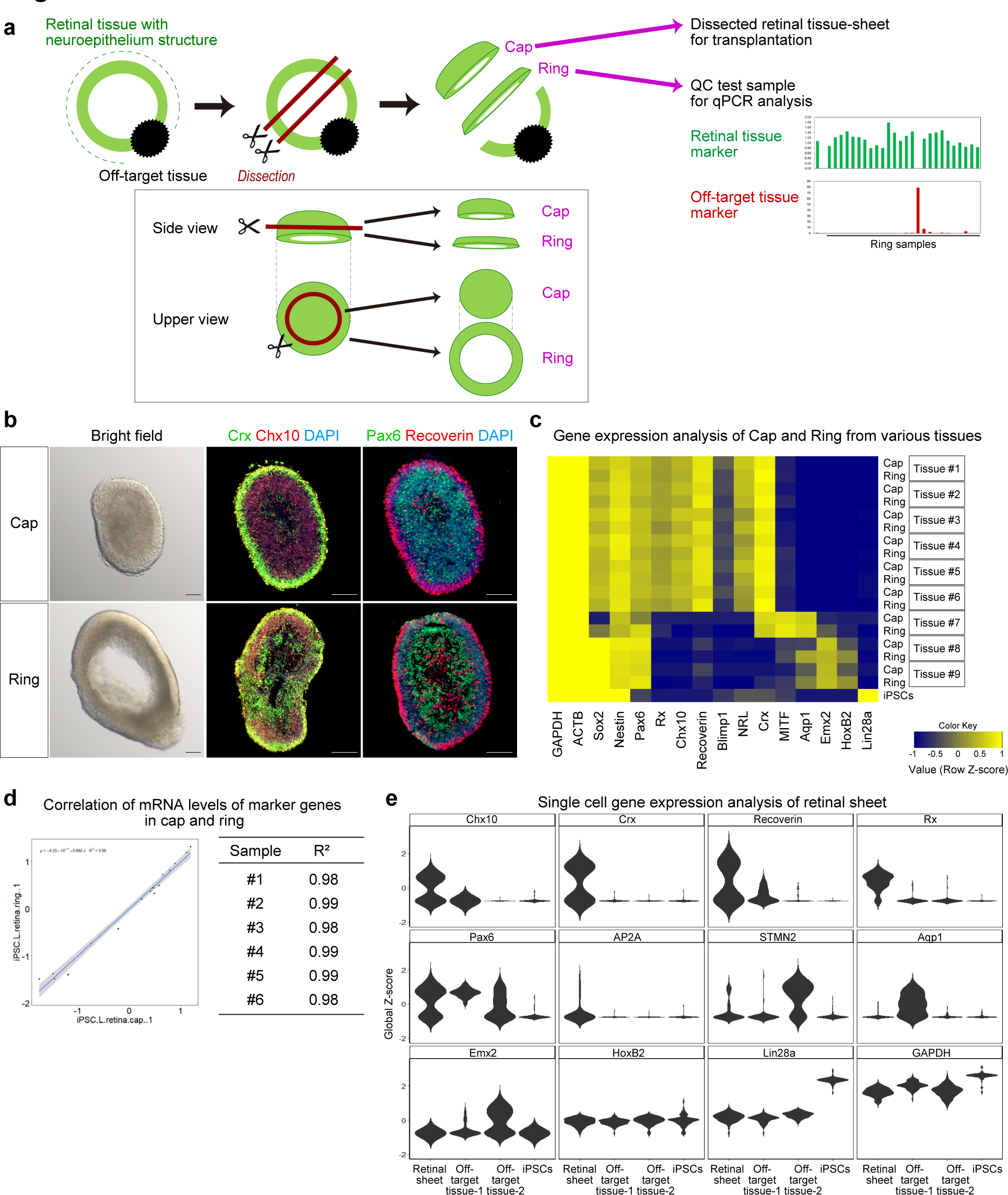
QC method for retinal sheets by testing gene expression in the surrounding outer tissue-sheet. **a** Scheme for dissection of the cap and ring and the ring-PCR test in Process 5 (dissection). **b** Bright-field (left) and IHC (middle, right) images of a representative cap and ring. The cap and ring were derived from iPSC-LPF11 cells. Immunostaining for Crx (middle, green), Chx10 (middle, red), Pax6 (right, green), and Recoverin (right, red) is shown. DAPI is shown in blue. Scale bar: 100 µm. **c** Heat map for gene expressions in caps and rings derived from iPSC-LPF11 cells. As a control, gene expressions in undifferentiated iPSC-LPF11 cells are also shown. Gene expressions were measured by qPCR analyses. Row Z-scores were calculated for each cap and ring, and plotted as a heat map. **d** Representative gene expressions in cap and ring dissected from the same aggregate shown as a plot with a regression line (left). The coefficient of determination (R^2^) for the gene expression between each cap and ring is shown (right). #1–#6 correspond to Tissue #1–#6 in (**c**). **e** Violin plots of single cell PCR data obtained from a retinal sheet, off-target tissue-1, off-target tissue-2, and undifferentiated iPSCs. Expressions of genes for photoreceptor cells (*Crx*, *Recoverin*), retinal progenitor cells (*Chx10*, *Rx*, *Pax6*), other retinal neurons (*Pax6*, *AP2A*, *STMN2*), and off-target tissues (*Aqp1*, *Emx2*, *HoxB2*, *Lin28a*) are shown.

The continuous neuroepithelium, self-formed in iPSC-derived 3D-retina, was dissected into the cap and ring (Fig. 3a, b). The cap and ring were both found to contain a multilayered continuous retinal tissue comprising a photoreceptor precursor layer with Crx^+^ and Recoverin^+^ cells on the outer surface and a retinal progenitor cell layer with Chx10^+^ and Pax6^+^ cells on the inner side.

We compared the gene expression patterns in the caps and rings. Various neuroepithelia were dissected into cap and ring tissues, and gene expression was determined by qPCR (Fig. 3c). We found that the gene expression patterns in each cap and ring from the same neuroepithelium were similar. To statistically analyze the similarity, we drew regression lines for the gene expression in each cap and ring and calculated the coefficient of determination (R^2^). We found that the R^2^ values for all sets of cap and ring were almost 1.0 (Fig. 3d and Supplementary Fig. 3a, b). These data indicate that the gene expression levels of retinal genes and off-target tissue-marker genes are almost same between the cap and ring dissected from the same aggregate.

Then, we carry out qPCR of each ring (Fig. 3a, ring-PCR test). If a ring expresses retinal genes and does not so much express off-target tissue marker genes such as tissue #1–6 in Fig. 3c, corresponding cap can be selected as the retinal sheet (final product) for transplantation. The retinal sheets, derived from four iPSC lines and passed ring-PCR test, indeed showed similar retinal gene expression patterns (Supplementary Fig. 3c). We further performed single cell PCR (scPCR) analysis and found that the majority of cells in retinal sheets expressed *Chx10*, *Crx*, *Recoverin*, *Rx*, *Pax6*, *AP2A*, or *STMN2* (Fig. 3e).

These results demonstrated that the ring-PCR test can estimate the gene expression in the cap to select retinal sheets comprised of retinal cells.

### Development of a controlled room temperature non-freezing preservation method

Between the dissection process and retinal sheet transplantation, it is necessary to perform QC tests including the ring-PCR test to select transplantable retinal sheets.

However, it requires at least 1–2 days to complete the ring-PCR test. Although retinal tissue can be frozen and thawed as reported previously^6, 20^, subretinal transplantation of frozen retinal tissue in nude rats showed impaired engraftment. We therefore aimed to develop a non-freezing preservation method for retinal tissue.

We first examined the optimal medium and temperature for non-freezing preservation of iPSC-derived 3D-retinas and dissected retinal sheets. To determine the optimal medium, we evaluated retinal maturation medium, University of Wisconsin (UW) solution, balanced salt solution (BSS), and Optisol-GS (Optisol hereafter), as candidate preservation solutions^49–51^, at 4, 17, and 37 °C. iPSC-derived 3D-retinas were generated, preserved for 3 days under various conditions, and cultured in retinal maturation medium at 37 °C for 7 days as recovery culture (Fig. 4a, b). The preserved retinas and control retinas were fixed and immunostained to check the morphology of the retinal tissue. We found that preservation at 17 °C with BSS or Optisol were optimal conditions to maintain the multilayered continuous retinal epithelium structure (Fig. 4a). Preservation at 4 °C was not a preferred condition, possibly causing damage to the retina. UW solution, a golden standard for preserving peripheral tissues including the pancreas, was also not preferred. One of the differences between UW solution and Optisol was the concentration of potassium chloride (KCl): 120 mM in UW solution and 5.33 mM in Optisol. Indeed, Optisol supplemented with KCl at 120 mM impaired the morphology of the retinal tissue (Supplementary Fig. 4a). Consistently, the concentration of KCl in BSS and Dulbecco’s modified eagle medium (DMEM, basal medium used for mammalian cell culture) was 5.3 mM and preservation at 17 °C in BSS and DMEM was found to be preferable (Fig. 4a and Supplementary Fig. 4b). These results raise the possibility that the concentration of KCl might be a key factor for the preservation solution. Based on these results, we chose Optisol for the non-freezing preservation solution, because Optisol is widely clinically used for human corneal transplantation^51^.

**Fig. 4.**
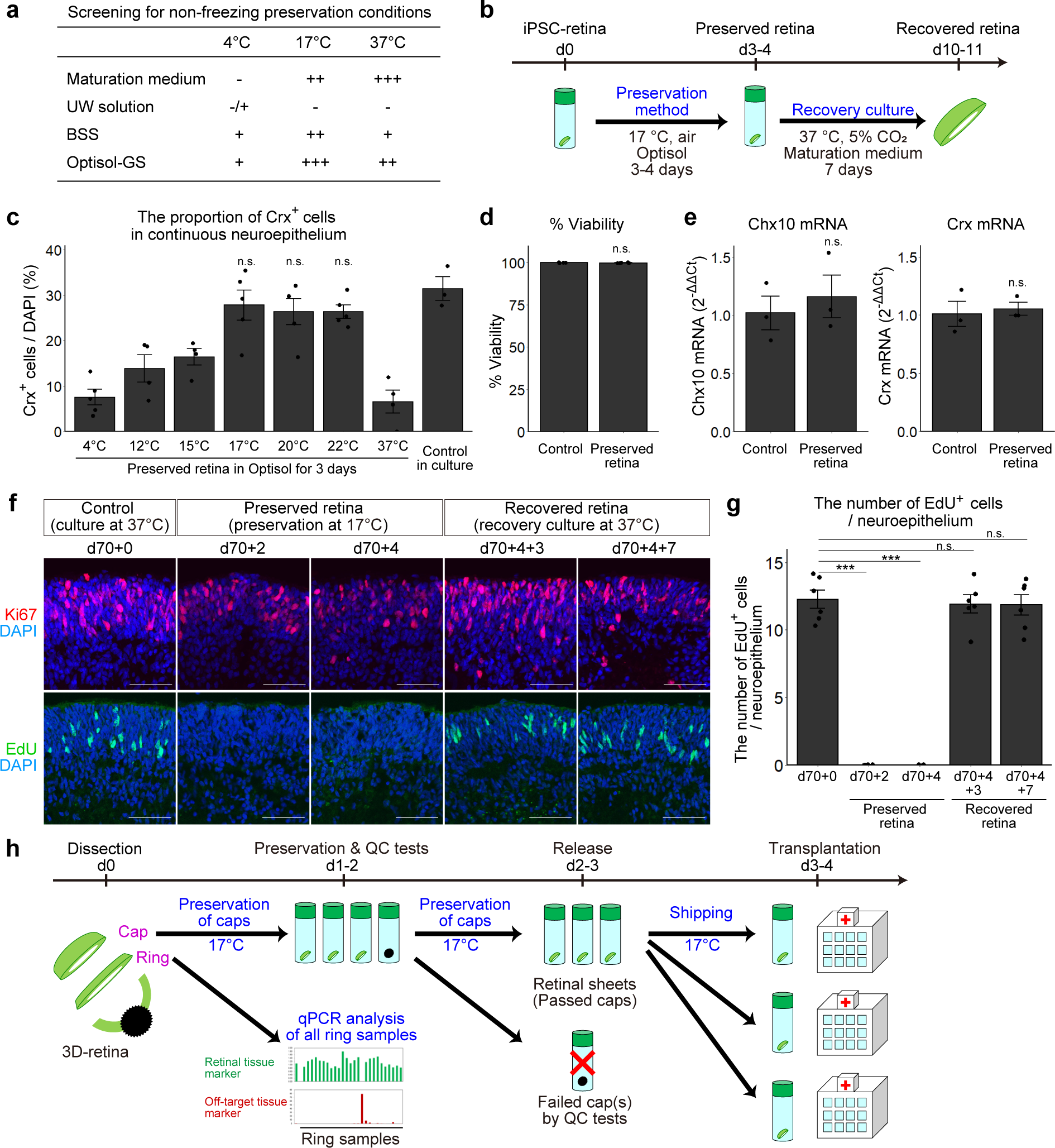
Controlled room temperature non-freezing preservation method. **a** Screening for the non-freezing preservation conditions. +++: multilayered continuous retinal epithelium was preserved. ++: multilayered continuous retinal epithelium was preserved but neural rosettes appeared. +: multilayered continuous retinal epithelium was observed but very limited. –, no multilayered continuous retinal epithelium was observed after the preservation. **b** Scheme of the preservation experiments. **c** Proportions of Crx^+^ photoreceptor precursors in multilayered retinal tissues. The proportions were calculated by dividing the number of Crx^+^ cells by the number of nuclei (DAPI). Data are presented as mean ± SE for *n* = 5 (4 °C), 4 (12 °C), 4 (15 °C), 5 (17 °C), 5 (20 °C), 5 (22 °C), 4 (37 °C), and 3 (control culture). **d** Cell viability in non-preserved retinas (Control) and retinas preserved for 4 days. Data are shown as mean ± SE (*n* = 5 per group). **e** Gene expressions of *Chx10* and *Crx* in non-preserved retinas (Control) and retinas preserved for 4 days. Data are presented as mean ± SE (*n* = 3 per group). **f** IHC analysis of retinas that were not preserved (d70+0), preserved in Optisol for 2 days (d70+2) and 4 days (d70+4), and preserved in Optisol for 4 days followed by recovery culture for 3 days (d70+4+3) and 7 days (d70+4+7). Ki67 (red) staining in the upper panels. EdU staining (green) in lower panels. DAPI is shown in blue. Scale bar: 50 µm. **g** Numbers of EdU-positive cells per 100 µm-wide multilayered retinal tissue in each group. Data are presented as mean ± SE (*n* = 6 per group). \*\*\**p*<0.001 for one-way ANOVA followed by a Tukey’s test. **h** Scheme for dissection, ring-PCR test, and shipping of retinal sheets. n.s., not significant.

To determine the optimal temperature for preservation, 3D-retinas were preserved in Optisol at 4, 12, 15, 17, 20, 22, and 37 °C for 3 days and then cultured in retinal maturation medium at 37 °C for 7 days as the recovery culture (Fig. 4b). We performed IHC analysis and found that the proportions of Crx^+^ cells were higher at 12,5, 17, 20, and 22 °C than at 4 and 37 °C and that the proportions of Crx^+^ cells at 17, 20, and 22 °C were much higher (Fig. 4c). Thus, the preferable range of preservation temperature was 17±5 °C and the optimal temperature was 17–22 °C.

We examined the cell viability and gene expression in retinas preserved in Optisol at 17 °C for 4 days. The cell viability in the preserved retinas exceeded 95% and was similar to that in control retinas in culture (Fig. 4d). The mRNA levels of *Chx10* and *Crx* in the preserved retinas were comparable to those in control retinas in culture (Fig. 4e). Furthermore, the mRNA levels of *Chx10* and *Crx* in retinas preserved in Optisol at 17 °C for 7 days were comparable to those in control retinas in culture. These results demonstrated that non-freezing preservation in Optisol at 17 °C was the optimal condition for retinal tissue.

We analyzed the status of retinal tissues preserved at 17 °C. Control retinas before preservation (d70+0), retinas preserved in Optisol at 17 °C for 2 days (d70+2) and 4 days (d70+4), and retinas recovered by culture at 37 °C for 3 days (d70+4+3) and 7 days (d70+4+7) were treated with the thymidine analog 5-ethynyl-2’-deoxyuridine (EdU) for 4 h to label proliferating cells in S-phase of the cell cycle and subjected to histological analysis. Expression of Ki67 was detected in both control retinas before preservation and retinas preserved at 17 °C (Fig. 4f, d70+2 and d70+4). In contrast, EdU^+^ cells were not detected in retinas preserved at 17 °C (Fig. 4f, g, d70+2 and d70+4), indicating that preserved retinal progenitors were arrested before entering S-phase. Importantly, EdU uptake was upregulated during recovery culture, indicating that the proliferation ability was reversible (Fig. 4f, g and Supplementary Fig. 4c, d). The number of cleaved caspase-3^+^ cells (apoptotic cells) was slightly high in the preserved retinas, while the numbers in the recovered retinas (d70+4+3 and d70+4+7) were similar to that in control retinas (Supplementary Fig. 4c).

By combining the controlled room temperature non-freezing preservation method (RT-preservation method hereafter) with the ring-PCR test, we developed a QC method for the dissection of 3D-retinas (Process 5) as follows: 1) dissection of the cap and ring from each 3D-retina, 2) preservation of the caps at 17 °C by the RT-preservation method, 3) the ring-PCR test to select passed and failed rings, 4) exclusion of caps corresponding to failed rings to select the retinal sheets (final products), and 5) shipping of the retinal sheets for transplantation (Fig. 4h).

### *In vivo* tumorigenicity study by subretinal transplantation in nude rats

An *in vivo* tumorigenicity study of the retinal sheet was conducted by subretinal transplantation in immunodeficient nude rats. Retinal sheets were generated from iPSC-Q cells and stored by the RT-preservation method for 3–4 days. The immunodeficient nude rats were divided into intact (*n* = 8), sham surgery (*n* = 12), and transplanted groups (*n* = 21), and the rats in the latter group underwent subretinal transplantation of one retinal sheet in the subretinal space (Supplementary Fig. 5a–f). Each transplanted retinal sheet (graft) was properly located in the subretinal space as confirmed by fundus imaging and optical coherence tomography (OCT) analysis (Supplementary Fig. 5b).

The transplanted rats showed no abnormalities in their body weights, survival rates, and clinical signs compared with intact or sham-operated rats during the observation period of up to 78 weeks after transplantation (Fig. 5a, b).

**Fig. 5.**
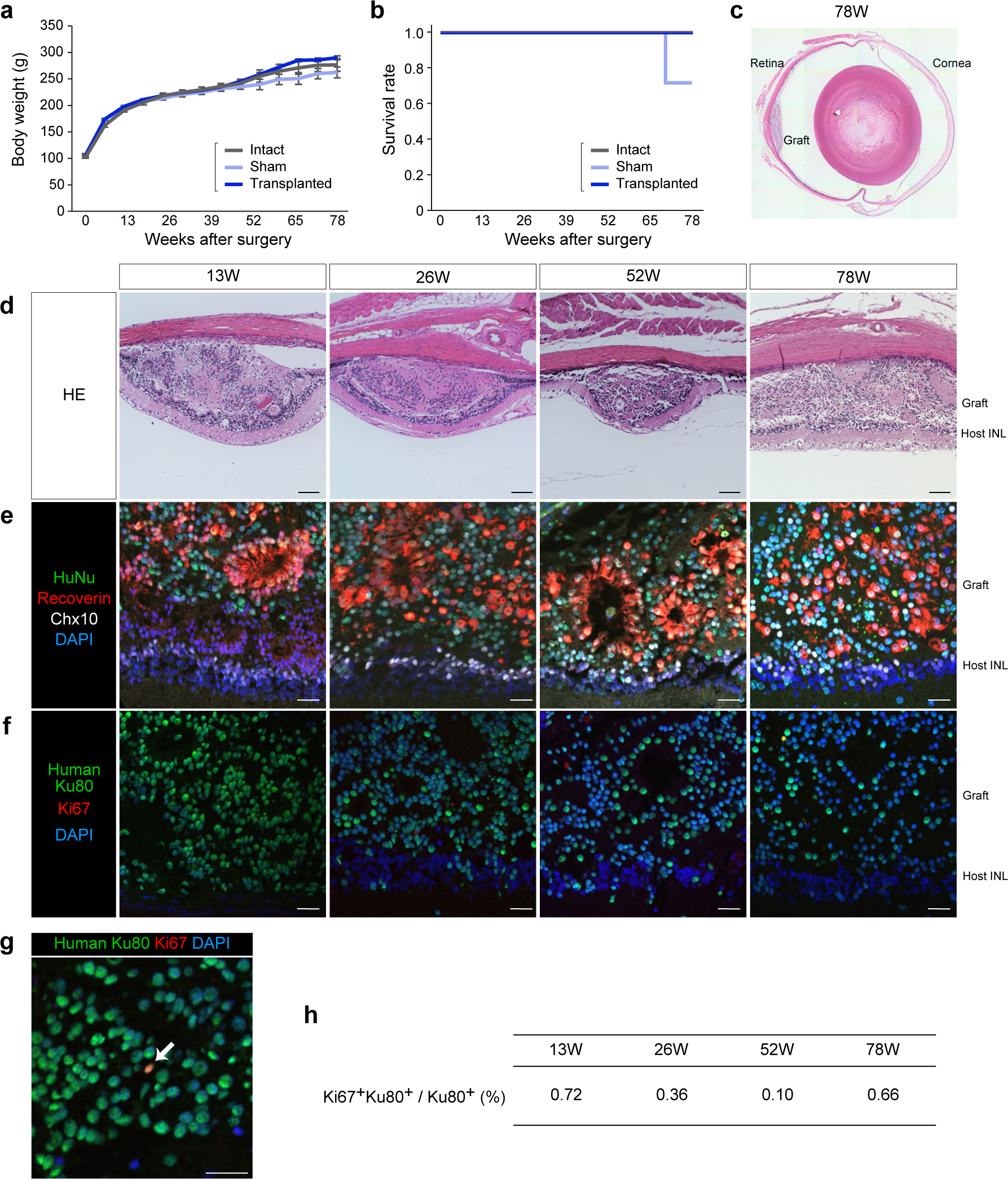
I*n vivo* tumorigenicity study by subretinal transplantation in nude rats. **a, b** Time courses of body weight (**a**) and Kaplan–Meier survival curves (**b**) in nude rats in the tumorigenicity study. Intact: control nude rat without surgery (*n* = 4). Sham: control nude rat with subretinal injection of vehicle control (*n* = 4). Transplanted: nude rat with subretinal transplantation of a single iPSC-Q-derived retinal sheet (*n* = 4). Data are presented as mean ± SE. **c, d** HE staining of transplanted eye sections at 13, 26, 52, and 78 weeks after transplantation. Scale bar: 100 µm. **e–h** Immunostaining of transplanted eye sections for human nuclear markers, retinal markers, and Ki67. Scale bar: 20 µm. (**e**) Immunostaining for HuNu (green), Recoverin (red), Chx10 (white), and DAPI (blue). (**f**) Immunostaining for human Ku80 (green), Ki67 (red), and DAPI (blue). (**g**) High-magnification image of immunostaining for the graft (52 weeks) with antibodies for human Ku80 (green), Ki67 (red), and DAPI (blue). Arrow indicates Ki67^+^ and Ku80^+^ cells. (**h**) Percentages of Ki67^+^ and Ku80^+^ cells among the Ku80^+^ human cells. Data are presented as mean. INL, inner nuclear layer.

Histopathological analysis of the grafts at 13, 26, 52, and 78 weeks after transplantation revealed no proliferative or malignant features such as nuclear atypia (Fig. 5c, d). IHC analyses of the grafts were then conducted (Fig. 5e–h). The percentages of proliferating human cells (Ki67^+^ and Ku80^+^) among the human cells (Ku80^+^) were <1% at each time point, indicating very limited proliferation of the graft cells (Fig. 5f, g, h)^52, 53^. Graft cells were found to differentiate into photoreceptors (Recoverin^+^ and HuNu^+^) and bipolar cells (Chx10^+^ and HuNu^+^) (Fig. 5e). Specifically, the iPSC-Q-derived retinal sheets became engrafted at >1.5 years after transplantation even under the xenotransplantation condition.

In this study, no tumorigenicity and no other adverse effects related to the iPSC-Q-derived retinal sheets were observed.

### Preclinical efficacy study by subretinal transplantation in retinal degeneration model rats

To examine whether preserved retinal sheets could become engrafted and differentiate into mature photoreceptors in severely degenerated condition, we performed a transplantation study using end-stage retinal degeneration model nude rats carrying a mutated human rhodopsin transgene (SD-Foxn1 Tg(S334ter)3Lav nude rats, RD-nude rats hereafter)^54, 55^. iPSC-derived retinal sheets stored by the RT-preservation method for 3–4 days were subretinally transplanted in RD-nude rats aged 20–30 weeks. At 24–44 weeks after transplantation, the PKCalpha^+^ and Recoverin^+^ host/rat bipolar cells still existed whereas the host/rat photoreceptors were almost completely degenerated at this stage (Fig. 6a and Supplementary Fig. 6a, control area). Recoverin^+^ and Ku80^+^ human photoreceptors became engrafted in the severely degenerated rat retinas to form photoreceptor rosette structures (Fig. 6a’ and Supplementary Fig. 6b, c, d), consistent with previous studies^8, 35, 36, 39^.

**Fig. 6.**
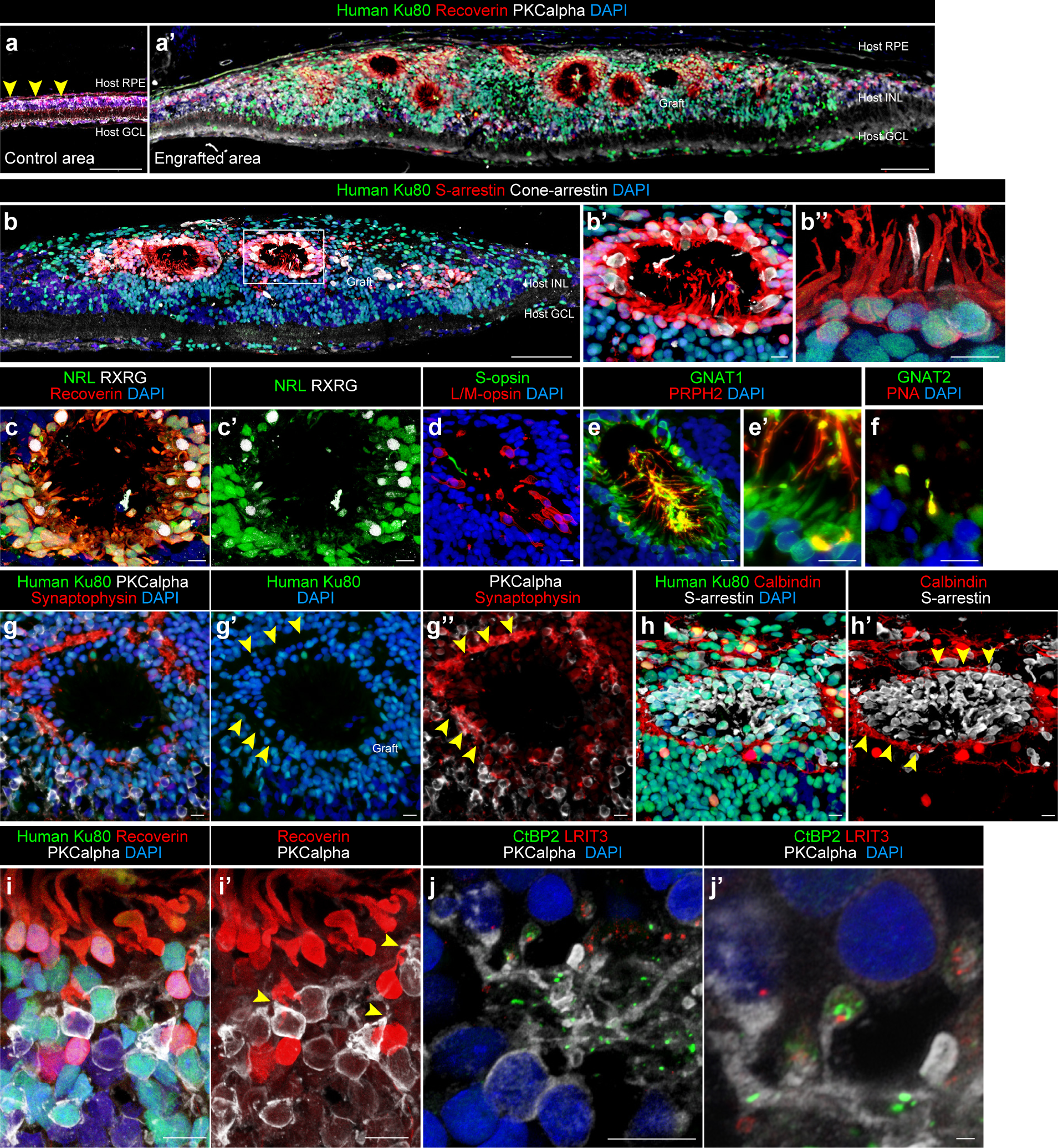
Engraftment and photoreceptor maturation of iPSC-retinal sheets after subretinal transplantation in RD-nude rats. **a–j’** Immunostaining of rat eyes transplanted with iPSC-S17-derived retinal sheets. The retinal sheets were transplanted in the subretinal space of RD-nude rats. The rat retinas were fixed at 263 days after transplantation (341 days after initiation of differentiation). **a, a’** Immunostaining for human Ku80 (green), Recoverin (red), and PKCalpha (white). Control non-transplanted area (**a**). Engrafted area (**a’**). Arrowheads in (**a**): human Ku80^−^, Recoverin^+^ and PKCalpha^+^ rat bipolar cells. **b–b’’** Immunostaining for human Ku80 (green), S-arrestin (red), and Cone-arrestin (white). Boxed area in (**b**) corresponds to (**b’**). High magnification in (**b’**) and higher magnification in (**b’’**). **c–h’** Immunostaining for photoreceptor markers. NRL (green), Recoverin (red), and RXRG (white) in (**c, c’**). S-opsin (green) and L/M-opsin (red) in (**d**). GNAT1 (green) and PRPH2 (red) in (**e, e’**). GNAT2 (green) and PNA (red) in (**f**). Human Ku80 (green), Synaptophysin (red), and PKCalpha (white) in (**g–g’’**). Arrowheads in (**g’, g’’**): Synaptophysin-positive neurites in no nuclear space. Human Ku80 (green), Calbindin (red), and S-arrestin (white) in (**h, h’**). Arrowheads in (**h’**): Calbindin-positive neurites. **i, i’** Maximum projection image of Z-stacks immunostained for human Ku80 (green), Recoverin (red), and PKCalpha (white) in (**i, i’**). Note that human Ku80^−^ and PKCalpha^+^ bipolar cells were located near the human Ku80^+^ and Recoverin^+^ photoreceptors (arrowheads). **j, j’** Maximum projection image of Z-stacks stained for CtBP2 (green), LRIT3 (red), and PKCalpha (white). High magnification in (**j’**). Note that photoreceptor-synapse marker CtBP2 and LRIT3 were expressed near the neurites of PKCalpha^+^ rod bipolar cells. DAPI staining (blue) in (**a, a’, b–b’’, c, d, e, e’, f, g, g’, h, i, j, j’**). Scale bars: 100 µm in (**a, a’, b**), 10 µm in (**b’, b’’, c–j**), and 1 µm in (**j’**). INL, inner nuclear layer; GCL, ganglion cell layer.

We further examined maturation of engrafted photoreceptors by the expression of rod, cone, outer segment (OS), outer plexiform layer (OPL), and synaptic markers. The photoreceptor rosettes contained both S-arrestin^+^ rod photoreceptors and Cone-arrestin^+^ cone photoreceptors (Fig. 6b, b’, b’’). The photoreceptor rosettes also contained NRL^+^ rod precursors, RXRG^+^ cone precursors, and some mature cone photoreceptors positive for S-opsin and L/M-opsin (Fig. 6c, c’, d). Rods and cones in the photoreceptor rosettes were positive for phototransduction markers (GNAT1 and GNAT2) and OS markers (PRPH2 and PNA), indicating that rods and cones maturated to form OS-like structures (Fig. 6e, e’, f). Synaptophysin^+^ neurites and Calbindin^+^ horizontal cells resided around the human photoreceptor rosettes, indicating that the OPL-like structure was formed (Fig. 6g, g’, g’’, h, h’, arrowheads). Higher magnification images showed that Recoverin^+^ and Ku80^+^ human photoreceptors were in direct contact with PKCalpha^+^ and Ku80^−^ host/rat bipolar cells (Fig. 6i, i’, arrowheads).

We also observed expression of the photoreceptor pre-synaptic markers CtBP2 (Ribeye) and LRIT3 at the tips of PKCalpha bipolar dendrites (Fig. 6j, j’). These results demonstrated that preserved retinal sheets became engrafted into severe RD-model immunodeficient rats to differentiate into mature photoreceptors with OS-like and OPL-like structures.

To demonstrate the light responsiveness of the retinal sheets, we performed *ex vivo* electrophysiology assays using multi-electrode array (MEA). iPSC-S17-derived retinal sheets on differentiation days 81–95 after RT-preservation for 3 days were transplanted in the subretinal space of 12 RD-nude rats aged 20–30 weeks.

Non-transplanted eyes were used as age-matched controls. The rats were subjected to *ex vivo* MEA analysis at 8–9 months after transplantation to evaluate the retinal ganglion cell (RGC)-derived spike counts and the RGC light responses as described previously^36, 56^. Light stimuli at three intensities (weak, medium, strong) were applied, and electrophysiological recordings were performed under three conditions: before addition of metabotropic glutamate receptor (mGluR6) blocker L-AP4 (Before), after addition of L-AP4 (L-AP4), and after washout of L-AP4 (After washout) (Fig. 7a, b). The length of engrafted area was approximately 1.0–2.0 mm (Fig. 7b, red line). The engrafted area was mounted on the electrodes. The peri-stimulus time histograms of the RGC-spike counts suggested that light stimulus-induced RGC-spikes were detected in the transplanted retina (Fig. 7c, upper). In a representative recording (Fig. 7c, box and Fig. 7d), increased RGC-spike counts were observed after light stimulus onset (Fig. 7d, Before). L-AP4 addition attenuated the light stimulus-induced RGC-spikes (Fig. 7c, d, L-AP4). After washout of L-AP4, light stimulus-induced RGC-spikes reappeared with enhanced responsiveness (Fig. 7d, After washout). In contrast, few light stimulus-induced RGC-spikes were detected in the non-transplanted control retinas (Fig. 7c, d, bottom). These results indicated that the RGC light responses in the transplanted rat retinas were derived from the grafted human retinal sheets and were dependent on synaptic transmission from grafted photoreceptors to bipolar cells. Similar RGC light responses were evident in 2 of 13 eyes transplanted with iPSC-Q-derived retinal sheets after RT-preservation (Supplementary Fig. 7a, b). Even iPSC-retinal sheets transplanted after RT-preservation for 4 days showed RGC light responses (and Supplementary Fig. 7c, d).

**Fig. 7.**
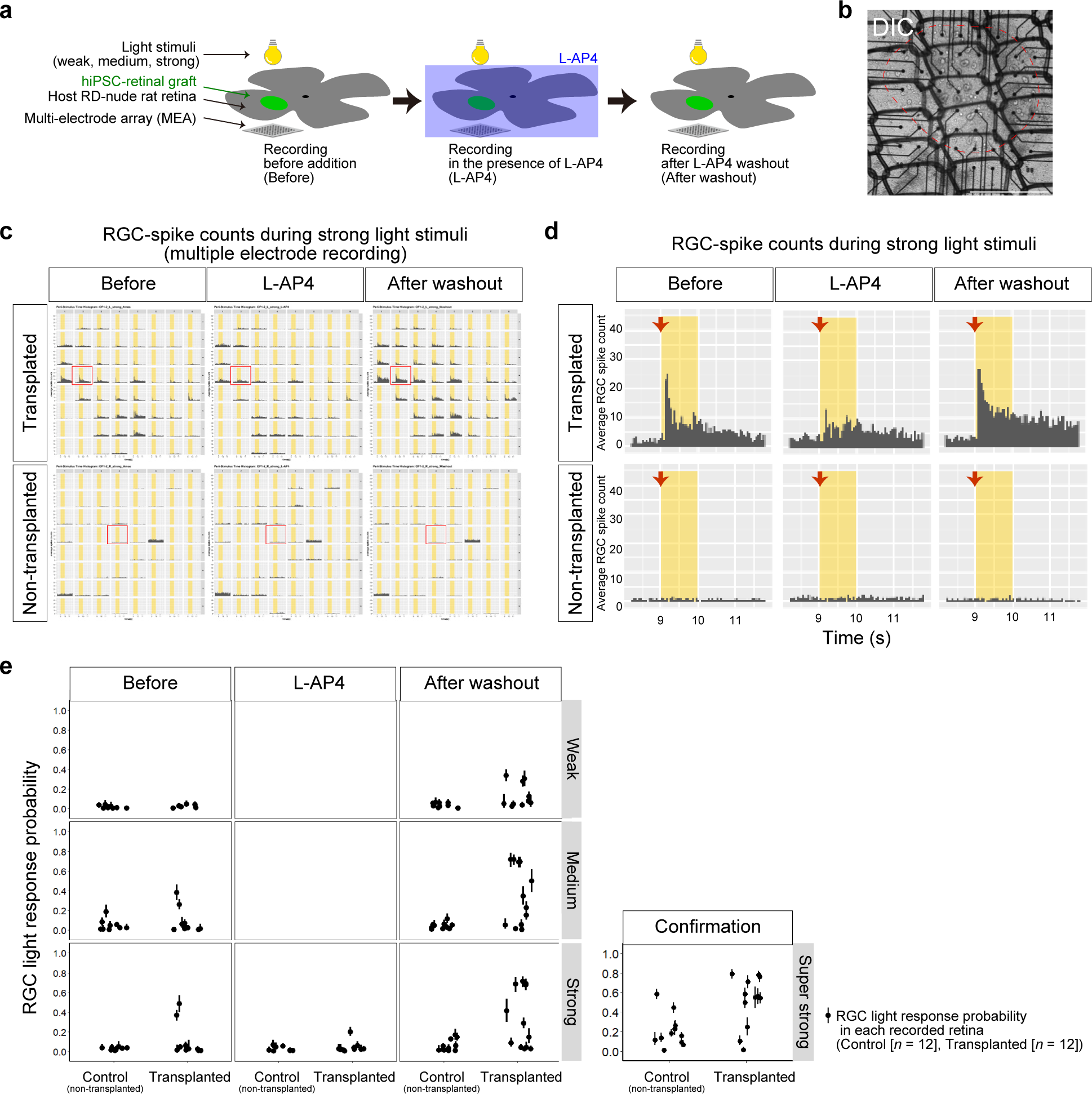
Light responses in transplanted iPSC-retinas by *ex vivo* electrophysiology assay. **a** Scheme of *ex vivo* electrophysiology assay using MEA to evaluate the light responses in transplanted iPSC-retinas. iPSC-S17-derived retinal sheets were transplanted in the subretinal space in RD-nude rats. Transplanted and non-transplanted rat retinas were mounted on the MEA electrodes for electrophysiological recording. During the recording, light stimuli were conducted in different intensities (weak, medium, strong), and the rat retinas were incubated in Ames’ medium (Before), followed by addition of L-AP4 (L-AP4), and washout of L-AP4 (After washout). **b–d** Representative results of MEA analysis. (**b**) Differential interference contrast (DIC) image of a RD-nude rat retina mounted on the MEA. Engrafted area was indicated by red dotted line. Scale bar: 1 mm. (**c, d**) Peri-stimulus time histograms to examine the RGC-spike counts during strong light stimuli (12.84 log photons/cm^2^/s). The X-axis and Y-axis represent the time and average RGC-spike count, respectively. Recording data for iPSC-S17-derived retinal sheet-transplanted rat retinas (Transplanted; upper) and non-transplanted rat retinas (Non-transplanted; lower) in the three conditions (Before, L-AP4, After washout) are shown. Multiple electrode recording data are shown in (**c**) and representative data (red box in **c**) are shown in (**d**). Yellow box in (**c, d**): light stimuli. Red arrows in (**d**): start timing of light stimuli. **e** RGC light response probabilities. Control non-transplanted rat retinas (Control; *n* = 12) and iPSC-S17-derived retinal sheet-transplanted rat retinas (Transplanted; *n* = 12) were subjected to MEA analysis to evaluate the RGC responses to light stimulation. Dots and bars show the per sample (recorded rat retina) summary of the collected data. Dot: RGC light response probability estimated by statistical modeling (Bayesian statistical inference with Markov Chain Monte Carlo sampling). Bar: 89% confidence interval.

To further confirm the functional potential of the retinal sheets, we compared the probability of RGC light responses in non-transplanted (Control) and transplanted retinas. The RGC number with light-responsive spikes per the number of recorded RGC (RGC light response probability) was estimated by using Bayesian statistical inference as described^56^. In non-transplanted rat retinas, robust RGC light responses were rarely detected for the weak, medium, and strong light stimuli (Fig. 7e, Control and Before; 0 light responses in 12 rat retinas, 0%). In rat retinas transplanted with iPSC-S17-derived retinal sheets, robust RGC light responses with average response probability > 0.25 were observed in two rat retinas for medium-to-strong light stimuli (Fig. 7e, Transplanted and Before; 2 light responses in 12 rat retinas, 17%). The detected RGC light responses were dominantly ON type because they showed attenuation in the presence of L-AP4 (Fig. 7e, Transplanted and L-AP4) and reappeared after L-AP4 was washed out (Fig. 7e, Transplanted and After washout). Note that RGC light-responses were enhanced after L-AP4 washout (Fig. 7e, Transplanted and After washout versus Transplanted and Before), consistent with the previous study^56^. After L-AP4 washout, RGC light responses were observed in three transplanted retinas for weak light stimulus and six transplanted retinas for medium-to-strong light stimuli (Fig. 7e, Transplanted and After washout; 6 light responses in 12 rat retinas, 50%). These results demonstrated that iPSC-retinal sheets transplanted after RT-preservation exhibited RGC light response functions (6 light responses in 12 transplanted rat retinas [50%] versus 0 light responses in 12 control rat retinas [0%]) (Fig. 7e).

Interestingly, RGC light responses were observed in retinal sheets transplanted on differentiation days 81–83 (*n* = 7) and 90–95 (*n* = 5), indicating that retinal sheets at approximately day 80 and approximately day 90 both had the potential to improve light responses in retinal degeneration model rats and that the allowable range of differentiation days for the efficacy may be wide (Supplementary Fig. 8a). In addition, RGC light responses in transplanted eyes were observed in both male and female RD-nude rats, indicating that retinal sheet transplantation enhanced RGC light responses in both male and female animals (Supplementary Fig. 8b).

Taken together, the efficacy studies demonstrated that iPSC-derived retinal sheets preserved at 17 °C for 3–4 days had the potential to become engrafted and mature in the severely degenerated RD-nude rat retinas to exhibit the RGC light response functions measured by *ex vivo* electrophysiology assays. These safety and efficacy studies on the retinal sheets demonstrated that the QC and RT-preservation methods were reasonable for the generation of transplantable retinal sheets (Supplementary Fig. 9).

## Discussion

Self-organizing stem cell culture technology enables the generation of 3D-tissues and organoids in a dish^1^. Self-organized tissue/organoid transplantation is an attractive approach for regenerative medicine in retinal degeneration. Previous studies have shown that hPSC-retinal sheets were engrafted for 0.5–2 years in the end stage photoreceptor degeneration model rodents and non-human primates^8, 35–37, 39, 57^. The key remaining technologies for tissue/organoid transplantation were a QC strategy and a preservation method. In this study, we developed a QC strategy for hPSC-derived retinal sheets. Retinal tissue in culture self-forms a unique morphological structure that is easy to distinguish under a microscope, and therefore retinal sheets can be dissected. To exclude possible errors in the selection of retinal tissue and avoidance of off-target tissues, we characterized the major off-target tissues and developed the ring-PCR test. Retinal tissue with its large continuous neuroepithelial structure can be dissected into two tissue-sheets: the inner-central sheet (cap) and the outer-peripheral sheet (ring) with similar gene expression patterns. Thus, qPCR analysis of the ring can be used to estimate the gene expression in the corresponding cap. As the ring-PCR test requires 1–2 days to complete, we developed the RT-preservation method to store retinal sheets for at least 3–4 days. To demonstrate the rationale for the QC strategy and the

RT-preservation method, we performed *in vivo* tumorigenicity and preclinical efficacy studies. In the tumorigenicity study, no tumors or no other adverse effects related to the iPSC-derived retinal sheets were observed. The preclinical efficacy study demonstrated that preserved retinal sheets had the potential to become engrafted and mature in the severely degenerated condition and exhibited light response functions measured by *ex vivo* electrophysiology assays. These preclinical studies support the concept of iPSC-derived retinal-sheet transplantation for retinal degeneration.

Based on present and previous studies^5–8, 27, 33–39^, Kobe City Eye Hospital applied for clinical research for a limited number of patients using iPSC-derived retinal sheets for retinitis pigmentosa. In June 2020, the clinical research plan was approved by the Health Science Council at the Ministry of Health, Labour and Welfare of Japan, and the clinical research was started (Trial ID: jRCTa050200027). Sumitomo Pharma developed a clinical grade manufacturing process and generated clinical grade allogeneic iPSC-derived retinal sheets. iPSC-retinal sheets were transplanted in two patients in 2020–2021.

The present study is based on a highly efficient and robust self-organizing culture technology. The combination of SFEBq, BMP, preconditioning, and induction-reversal methods allows robust self-organizing 3D-differentiation culture for retinal tissue generation with a large continuous neuroepithelium^7, 8^. This large continuous neuroepithelium structure had a size of 1–3 mm in the tangential direction and was easy to distinguish under the microscope, making it possible to dissect the cap and ring. The self-organized retinal tissue with continuous neuroepithelium reproducibly differentiated to form a multilayered structure with Crx^+^ photoreceptor precursor and Chx10^+^ retinal progenitor cell layers from multiple hPSC lines using our protocols or different culture protocols^6, 12–14, 18, 19, 22^. This feature of 3D self-organization possibly based on the intrinsic embryonic development program is the core technology for robust generation of retinal sheets with controlled quality.

One of the features of the QC method in the present study was the dissection and analysis of the ring to check the quality of the cap. This cap-and-ring approach could be utilized for other organoids with a continuous epithelium such as cortical, hippocampal, pituitary, and intestinal organoids^40, 58–60^. Although manual dissection of tiny tissues is a prevalent experimental procedure in research using Xenopus early embryos or mouse developing brains for primary neural precursor cell culture^33, 39^, automation and mechanization of the dissection process will be important for future manufacture of iPSC-derived retinal sheets.

To examine the quality of the ring, one candidate QC test was qPCR analysis and the others were IHC analysis, flow cytometry, gene chip analysis (microarray), RNA-seq, and single cell analysis. We chose qPCR in this study, because it allows high-throughput assays to test many samples in a relatively short period, has high sensitivity, and is technically easy^48^. The other QC tests also had strong points. A recent study using RNA-seq approach revealed that Six6-negative off-target cells in retinal differentiation culture using Matrigel method contain multiple neural cells including Emx2-positive cells and HoxB2-positive cells^61^. In addition, single cell analyses such as single cell RNA-seq are powerful technologies, and several groups have reported pioneered studies^13, 62–65^. Meanwhile, imaging technologies and computational image processing, such as deep-learning technologies, are promising approaches for the development of non-destructive QC tests. In future studies, we would like to proceed with single cell analysis and imaging technology to further investigate the quality of the self-organizing 3D-retinas.

Although cryopreservation is an attractive approach for worldwide supply, cryopreserved retinal sheets exhibited impaired engraftment. Therefore, we needed to develop a non-freezing preservation method and found that the optimal temperature for retinal sheet preservation was nearly room temperature (17±5 °C). Room temperature preservation was reported for hepatic tissue^49^, and our study demonstrated that near room temperature was suitable for neural tissues like retinal tissue. Low temperatures around 4 °C caused damage to the tissues, possibly through abnormal intracellular metabolism at 4 °C^49^. Culture at 37 °C under 5% CO_2_ was suitable for cell growth, but the morphology of the retinal sheets changed during self-organizing culture.

RT-preservation at 17±5 °C could maintain the morphology of the retinal sheets. During RT-preservation, the retinal tissue arrested to enter S-phase, raising the possibility that proliferation may become paused under this preservation condition. RT-preservation was recently reported for RPE and retinal tissue^66, 67^. We developed an RT-preservation method and performed transplantation studies, including a 1.5-year tumorigenicity study, for retinal sheets under the RT-preservation condition.

Toward clinical applications, we evaluated the *in vivo* tumorigenicity and efficacy of iPSC-retinal sheets generated using the ring-PCR test and RT-preservation method. In tumorigenicity study, no tumors or no other adverse effects related to the retinal sheets were observed. RT-preserved iPSC-retinal sheets became engrafted and matured in RD-nude rats to form photoreceptor rosettes. Similar photoreceptor rosette structures were observed in human donor-derived fetal retinal tissue transplantation in rats^68^. Although a half of human iPSC-photoreceptors in rosettes on host RPE side were far from host/rat bipolar cells, the other half of human iPSC-photoreceptors in rosettes on host RGC side could contact to host/rat bipolar cells. Engrafted iPSC-photoreceptors *in vivo* matured to form the OS-like structure on the inner-apical surface of the rosettes and the OPL-like structure with photoreceptor synaptic marker expression in the outer basal side adjacent to bipolar cells and horizontal cells. Electron microscopy will be a key to demonstrate the synaptic connections between grafted photoreceptors and host bipolar cells. Importantly, 50% of iPSC-retinal transplants exhibited light responses monitored by *ex vivo* MEA recordings. This rate was comparable to previous results for transplantation of fresh iPSC-retinas without RT-preservation (4 light responses in 7 transplanted rat eyes, 57%)^36^. These observations demonstrated that RT-preserved iPSC-retinal sheets transplanted in the end stage retinal degeneration condition became engrafted and differentiated into mature photoreceptors and exhibited RGC light responses. In future study, it will be intriguing to investigate the light responsiveness of RT-preserved retinal sheets in the brain and behavior levels^37^.

Toward future clinical applications in large numbers of patients, further preclinical studies will be a key. Among them, the large animal studies to develop surgical procedures, transplantation devices, and immunosuppression protocols are important to enable widespread use of retinal sheets. Based on the self-organizing stem cell culture technology, the realization of tissue/organoid therapy in medicine is ongoing.

## Methods

### Human iPSC lines

Two human iPSC lines, iPSC-LPF11 and iPSC-S17, were established from peripheral blood cells using Sendai virus vectors by Sumitomo Pharma^8^. The iPSC-QHJI01s04 line (iPSC-Q) was established by the iPSC Stock Project organized by the Kyoto University Center for iPS Cell Research and Application (CiRA), and provided from Kyoto University^69, 70^. The iPSC-1231A3 line established at Kyoto University and derived from ePBMC(R) purchased from Cellular Technology Limited (http://www.immunospot.com/) was provided by Kyoto University^41^. The experimental protocols using clinical grade human iPSCs were approved by the Research Ethics Committee of Sumitomo Pharma Co., Ltd., Japan.

### Self-organizing culture of human iPSCs for 3D-retina generation

The feeder-free hPSC-culture, SFEBq, preconditioning, d0-SAG, BMP, and induction-reversal culture methods were performed as described^6–8, 41^ with slight modifications. In Process 1 (maintenance culture), human iPSCs were maintained on LM511-E8 matrix (Nippi) in StemFit medium (Ajinomoto) according to a published protocol^41^ with slight modifications^8^. The medium was changed every 1–2 days until the cells reached 70%–80% confluence. iPSCs were passaged using TrypLE Select Enzyme (Thermo Fisher Scientific) and dissociated into single cells by gentle pipetting. The dissociated iPSCs were seeded at a density of 700–1700 cells/cm^2^ and cultured in LM511-E8 matrix-coated six-well culture plates (Iwaki) containing StemFit medium with 10 µM Y-27632 (Wako)^8^. Prior to differentiation, iPSCs were treated with SB431542 (SB; Wako) and/or smoothened agonist (SAG; Enzo Biochem) as the preconditioning step. In Process 2 (retinal differentiation), hPSCs were treated with TrypLE Select Enzyme at 37 °C for 4–7 min, and dissociated into single cells by gentle pipetting. The dissociated iPSCs were quickly reaggregated using low-cell-adhesion 96-well plates with V-bottomed wells (Sumilon PrimeSurface plates; Sumitomo Bakelite) in differentiation medium (gfCDM+KSR) with Y-27632 and SAG (d0-SAG method). The differentiation medium was gfCDM supplemented with 10% KSR, while gfCDM alone comprised 45% Iscove’s modified Dulbecco’s medium (Gibco), 45% Hams F12 (Gibco), Glutamax, 1% Chemically Defined Lipid Concentrate (Gibco), and 450 µM monothioglycerol (Sigma-Aldrich). The day of SFEBq culture initiation was defined as day 0. On day 3, recombinant human BMP4 (R&D Systems) was added at 1.5 nM (55 ng/ml)^7^. Thereafter, the culture medium was changed every 3–4 days to generate immature retinal tissue.

In Process 3 (induction-reversal culture), the immature retinal tissue generated from iPSCs was subjected to a two-step induction-reversal culture as follows. For the induction culture, cell aggregates on days 14–18 were transferred from 96-well plates to 90-mm non-cell-adhesive petri dishes (Sumitomo Bakelite; approximately 32–48 aggregates/90-mm dish), and cultured for 3 days in DMEM/F12-Glutamax medium (Gibco) containing 1% N2 supplement (Gibco), 3 µM CHIR99021 (GSK3 inhibitor; Wako), and 5 µM SU5402 (FGFR inhibitor; Wako). For the reversal culture (Process 3) and the maturation culture (Process 4), the cell aggregates were cultured in retina maturation medium as described (Nukaya et al. WO2019017492A1, WO2019054514A1). The medium was changed every third or fourth day to obtain 3D-retinas. Floating cell aggregates were analyzed using an inverted microscope (Keyence BZ-X810, Nikon Eclipse-Ti, or Olympus IX83).

### Dissection of the cap and ring

Each cell aggregate containing continuous stratified neuroepithelial tissue was subjected to dissection of the cap and ring (Process 5). The central part of the neuroepithelial tissue was cut from the cell aggregate under a microscope as described^39^. The dissected inner-central neuroepithelial tissue sheet was collected as the cap. At the same time, the surrounding outer-peripheral part of the neuroepithelial tissue was cut from the cell aggregate and harvested as the ring. The obtained caps and rings were subjected to the RT-preservation method and the ring-qPCR test, respectively. Caps whose rings passed the ring-PCR test were used as retinal sheets for transplantation.

### qPCR analysis and ring-PCR test

Rings and other tissue samples were lysed with Buffer RLT (Qiagen) containing 1% 2-mercaptoethanol, and total RNA was extracted and purified using an RNeasy Micro Kit (Qiagen). The total RNA was reverse-transcribed and subjected to qPCR using a StepOne Plus Real-Time PCR System (Applied Biosystems) or Biomark HD (Fluidigm) according to the manufacturers’ instructions. Based on the Ct values for each gene, Z-scores were calculated for the individual caps or rings. Data were visualized as heat maps using the heatmap.2 function of the R/Bioconductor package gplots. PCA was performed using the prcomp function in the R base package. The PCA results were visualized using R package ggplot2. The qPCR primers (TaqMan probes) were purchased from Applied Biosystems, Inc. and used for qPCR analysis.

### Microarray analysis

Dissected tissues in the cell aggregates were lysed with Buffer RLT (Qiagen) containing 1% 2-mercaptoethanol, and total RNA was extracted and purified using an RNeasy Micro Kit (Qiagen). Microarray analysis using a GeneChip array was performed by Kurabo Industries (Osaka, Japan). Briefly, total RNA was reverse-transcribed to cDNA with T7 oligo d(T) primer (Affymetrix). Next, biotin-labeled cRNA was synthesized and amplified by *in vitro* transcription of the second-strand cDNA template using T7 RNA polymerase (Affymetrix). The labeled cRNA was purified and fragmented, loaded onto a GeneChip® Human Genome U133 Plus2.0 array (Affymetrix), and hybridized according to the manufacturer’s protocol. Raw intensity data from the GeneChip array were analyzed using GeneChip Operating Software (Affymetrix). The data were logarithmically transformed and logFC was calculated for each gene. The logFC data were visualized as heat maps using the heatmap.2 function of the R/Bioconductor package gplots.

### Single cell polymerase chain reaction (scPCR) analysis

Caps and rings were dissected from 3D-retinas. The cap whose ring passed the ring-PCR test was subjected to scPCR analysis. The cap was dissociated into single cells by using papain (Wako), followed by loaded into the C1 preamp IFC (10–17 μm, Fluidigm) for single cell isolation. Pre-amplification of genes was performed in the C1 instrument (Fluidigm) according to manufacturer’s instructions. The pre-amplified cDNAs from each cell were further amplified by using the Biomark HD instrument (Fluidigm) with the TaqMan assays.

### RT-preservation method

Cell aggregates and tissue sheets (caps) were transferred to 1.5-mL or 15-mL tubes. After washout of the culture supernatant, preservation solution was added and the tubes were placed in incubators set at 4, 12, 15, 17, 20, 22, and 37 °C. For temperature validation, the temperatures of the incubators were monitored by temperature loggers in some experiments. As candidate preservation solutions, Optisol-GS (Bausch & Lomb), balanced salt solution (BSS) (Gibco), Belzer UW Cold Storage Solution (Bridge to Life), and cell culture media were used. For RT-preservation of caps, each 1.5-mL tube containing a cap in Optisol was placed in a cool incubator (Mitsubishi Electric Engineering) set at 17 °C. Caps that passed QC tests, including the ring-PCR test were picked up and used for transplantation.

### Cell viability assay

Tissue sheets which were dissected from 3D-retinas were dissociated into single cells by using papain (Wako). After centrifugation and removal of supernatants, dissociated cells were suspended in fresh media. Total cells and dead cells were stained by acridine orange and DAPI, and the number and the percentage of viable cells were determined by using NucleoCounter® NC-200 (ChemoMetec).

### Animals, transplantation in rats, and *ex vivo* electrophysiology assay

All the animal experimental protocols were approved by the animal care committee of the RIKEN Center for Biosystems Dynamics Research (BDR) and were conducted in accordance with the Association for Research in Vision and Ophthalmology Statement for the Use of Animals in Ophthalmic and Vision Research. SD-Foxn1 Tg(S334ter)3Lav nude rats (RD-nude rats) were obtained from the Rat Resource and Research Center^54, 55^.

Transplantation in the subretinal space of rats was performed as described^36^. Both male and female RD-nude rats transplanted with iPSC-retinal sheets were used for MEA recordings at 13.5–15 months of age (i.e., 8–10.5 months after transplantation). The MEA recordings were performed using the USB-MEA60-Up-System (Multi Channel Systems) as described^36, 56^. Briefly, RD-nude rats were dark-adapted for 1–3 days before use. Rats were anesthetized and sacrificed by excess inhalation of Isoflurane or Sevoflurane. Their eyecups were then harvested under dim red light with wavelength peaked at 700 nm and placed in oxygenated Ames’ medium (Sigma-Aldrich) in the dark. The retina was carefully isolated from the eyecup and the residual vitreous was removed.

The retina was mounted with the RGC side down. The engrafted area was recognized by its dotted appearance and centered on the electrode. Full-field light stimuli with 10 ms and 1 sec duration at different intensities (weak: 10.56 log photons/cm^2^/s; medium: 12.16 log photons/cm^2^/s; strong: 12.84 log photons/cm^2^/s) were generated using a white LED (NSPW500C; Nichia Corp.). Each set of stimulation, composed of 3 repeats for each stimulation intensity and duration combination, was repeated before addition of L-AP4 (Before), in the presence of 10 μM L-AP4 (L-AP4, mGluR6 blocker, Wako), and after washout of L-AP4 (After washout). Super-strong stimuli (15.48 log photons/cm^2^/s) were applied at the end of the MEA recordings to confirm the cell viability in the retinas. The MEA data were collected at a 20-kHz sampling rate without filtering. The recorded spikes were sorted offline to count RGC-spikes using the automatic template formation and spike matching algorithm in Spike 2 (version 7.2; CED) with minor modifications^56^.

RGC light responses were defined as having 2-fold increase of spiking frequency against the stimulation onset and/or offset. The average RGC-spike counts and the average RGC light response probability (RGC number with light-responsive spikes per number of recorded RGC) was calculated from all detected RGCs in each sample as described^36, 56^.

### *In vivo* tumorigenicity study

*In vivo* tumorigenicity study using animals was approved by the IRB of the Foundation for Biomedical Research and Innovation (FBRI), the Committee for Animal Experiments of the FBRI, and the animal care committee of RIKEN BDR. Subretinal transplantation was conducted in RIKEN BDR. Female nude rats were used in this study to reduce fighting between rats bred in a same cage^71^.

Six-week-old female nude rats (F344/NJcl-*rnu/rnu* rats; CLEA Japan) were used for the transplantation and underwent general condition observations and weight measurements during the observation period. Surgical stress of subretinal transplantation in rats was evaluated in sham-operated eyes (Supplementary Fig. 5e,f). Fundus imaging and OCT analysis were performed using RS-3000 Advance (Nidek) and Micron IV and OCT (Phoenix research labs) according to the manufacturers’ instructions. After necropsy, the eyeballs were fixed at 4 °C in SUPERFIX (KY-500; Kurabo Japan), embedded in paraffin, and sliced with a microtome at 3-µm thickness. One in every five sections was used for HE staining and histopathological analysis, and the other sections were used for immunohistochemistry.

### Immunohistochemistry

IHC was performed as described^8^. Cell aggregates and transplanted retinas harvested from RD-nude rats were fixed with 4% paraformaldehyde (Wako) and sectioned with a cryostat (Leica) to prepare frozen sections. Frozen sections and paraffin sections were treated with or without heat-based antigen retrieval in Target Retrieval solution (Dako) at 105 °C for 15 min. The primary antibodies used were as follows: anti-Chx10 (sheep; 1:500; Exalpha), anti-Pax6 (mouse; 1:1000; BD Biosciences), anti-Rx (guinea pig; 1:2000; Takara), anti-Crx (rabbit; 1:200; Takara), anti-Recoverin (rabbit; 1:500; Proteintech), anti-RXRG (mouse; 1:500; Santa Cruz Biotechnology), anti-NRL (goat; 1:500; R&D Systems), anti-Lhx2 (rabbit; 1:500; Millipore), anti-Rhodopsin (mouse; 1:1000; Sigma Aldrich), anti-S-opsin (goat; 1:1000; Santa Cruz Biotechnology), anti-S-opsin (rabbit; 1:500; Santa Cruz Biotechnology), anti-L/M-opsin (rabbit; 1:500; Millipore), anti-Cone-arrestin (Arrestin-3; goat; 1:500; Novus), anti-S-arrestin (mouse; 1:500; Novus), anti-CtBP2 (mouse; 1:500; BD Biosciences), anti-PKCalpha (goat; 1:500; R&D Systems), anti-Ki67 (mouse; 1:500; BD Biosciences), anti-Aqp1 (Aquaporin1; rabbit; 1:500; Millipore), anti-Emx2 (sheep; 1:100; R&D Systems), anti-cleaved caspase-3 (rabbit; 1:200; Cell Signaling Technology), anti-HuNu (mouse; 1:500; Millipore), anti-human Ku80 (rabbit; 1:500; Cell Signaling Technology), anti-human Ku80 (goat; 1:500; R&D Systems), anti-Gnat1 (rabbit; 1:500; Santa Cruz Biotechnology), anti-Gnat2 (rabbit; 1:500; Santa Cruz Biotechnology), anti-PRPH2 (mouse; 1:500; Millipore), Peanut agglutinin (PNA) (1:500; Thermo Fisher Scientific), anti-Synaptophysin (goat; 1:500; R&D Systems), and anti-LRIT3 (rabbit; 1:500; Novus). Nuclear counterstaining was performed with DAPI (Nacalai). Stained sections were analyzed with a fluorescence microscope (Keyence BZ-X810) or a laser scanning confocal microscope (Olympus Fluoview FV1000D, Leica TCS SP-8, or Carl Zeiss LSM880). Image analyses were performed with IMARIS (Oxford Instruments), ImageJ (National Institutes of Health), and Zen Blue (Carl Zeiss) imaging software.

### EdU uptake analysis

For EdU labeling, control retinas before preservation (d70+0), retinas preserved in Optisol at 17 °C for 2 days (d70+2) and 4 days (d70+4), and retinas recovered in culture at 37 °C for 3 days (d70+4+3) and 7 days (d70+4+7) were treated with the thymidine analog EdU for 4 h. The EdU-treated retinas were fixed and sectioned with a cryostat (Leica). Sections of the retinas were stained with Alexa Fluor 488-labeled azide (Invitrogen) according to the manufacturer’s protocol. Images were obtained using a confocal laser microscope and the numbers of EdU-positive cells and DAPI-positive nuclei were counted using ImageJ software (National Institutes of Health).

### Statistics and reproducibility

Statistical analyses were performed with R version 3.6.0 (The R Foundation for Statistical Computing). A two-tailed Student’s *t*-test was carried out for two-group comparisons, and one-way analysis of variance (ANOVA) followed by Tukey’s test was performed for multiple-group comparisons. Intergroup differences in body weight and survival rate were tested by one-way ANOVA and the log-rank test, respectively.

### Data availability

All reasonable requests will be promptly reviewed by the senior authors to determine whether the request is subject to any intellectual property or confidentiality obligations. This study did not generate new cell lines.

## Acknowledgements

We are grateful to the CiRA (Kyoto, Japan) and CiRA Foundation (Kyoto, Japan) for kindly providing iPSCs. We thank Dr. Mototsugu Eiraku for advice on retinal maturation culture, and Yasushi Hiramine, Miki Iwata, Kazunari Tanaka, Masahiro Yahata, Koichiro Manabe, Kiyoko Bando, Jiro Akimaru, Keigo Kawabe, Kenji Yoshida, Satoshi Ando, Atsushi Tsuchida, Tomokazu Nagano, and members of RACMO for fruitful discussions. We thank Ayumi Kiso for in vivo studies, Kohei Kanata for IHC staining, and Yukiko Ishigami for dissection. A.Ku. would like to express deep gratitude to his late mentor Dr. Yoshiki Sasai, a gifted scientist who pioneered self-organizing stem cell biology. This work was supported by the Research Center Network for Realization of Regenerative Medicine from the Japan Agency for Medical Research and Development (AMED) (M.T., S.K.).

## Author contribution

K.W. designed the study for off-target tissue analysis, ring-PCR test, and RT-preservation method, performed experiments, analyzed data, and wrote the manuscript. S.Y. designed the study for self-organizing culture and IHC analysis in vivo, performed experiments, analyzed data, and wrote the manuscript. H.Y.T. designed and performed electrophysiology assay, analyzed data, and wrote the manuscript. M.S., T.Ka., H.A., T.T., Y.K. designed and performed tumorigenicity study, analyzed data, and wrote the manuscript. A.N., K.U., K.O., C.M., T.M., J.S., M.N., O.T., O.O., M.F., Y.H., A.T., R.H., H.O., T.H., D.N., K.M. performed experiments and analyzed data. A.I., M.T., A.Ki., T.Ki. managed the project and analyzed data. S.K. supervised and designed in vivo tumorigenicity study, analyzed data, and wrote the manuscript. M.M. conceived, supervised and designed in vivo efficacy study, performed experiments, analyzed data, and wrote the manuscript. A.Ku. conceived, supervised and designed self-organizing culture, off-target tissue analysis, ring-PCR test, and RT-preservation method, performed experiments, analyzed data, wrote the manuscript, and made final approval of the manuscript.

