## Supplementary Material for "Self-organization, quality control, and preclinical studies of human iPSC-derived retinal sheets for tissue-transplantation therapy"

#### Supplementary Figures

**Supplementary Fig 1.** Self-organizing culture of human iPSCs to generate the 3D-retina and dissected retinal sheet (supplementary to Fig. 1).

**Supplementary Fig 2.** Characterization of major off-target tissues in the retinal differentiation culture (supplementary to Fig. 2).

**Supplementary Fig 3.** QC method for retinal sheets by testing the gene expression in the surrounding outer tissue-sheet (supplementary to Fig. 3).

**Supplementary Fig 4.** Controlled room temperature non-freezing preservation method (supplementary to Fig. 4).

**Supplementary Fig 5.** *In vivo* tumorigenicity study by subretinal transplantation in nude rats (supplementary to Fig. 5).

**Supplementary Fig 6.** Engraftment and photoreceptor maturation of retinal sheets after subretinal transplantation in RD-nude rats (supplementary to Fig. 6).

**Supplementary Fig 7.** Light responses in transplanted iPSC-retinas by the

MEA assay (supplementary to Fig. 7).

**Supplementary Fig 8.** RGC light response probabilities in transplanted iPSC-retinas by the MEA assay (supplementary to Fig. 7).

**Supplementary Fig 9.** Five human PSC lines and their preclinical data in the previous and present studies.

#### Supplementary Figure 1

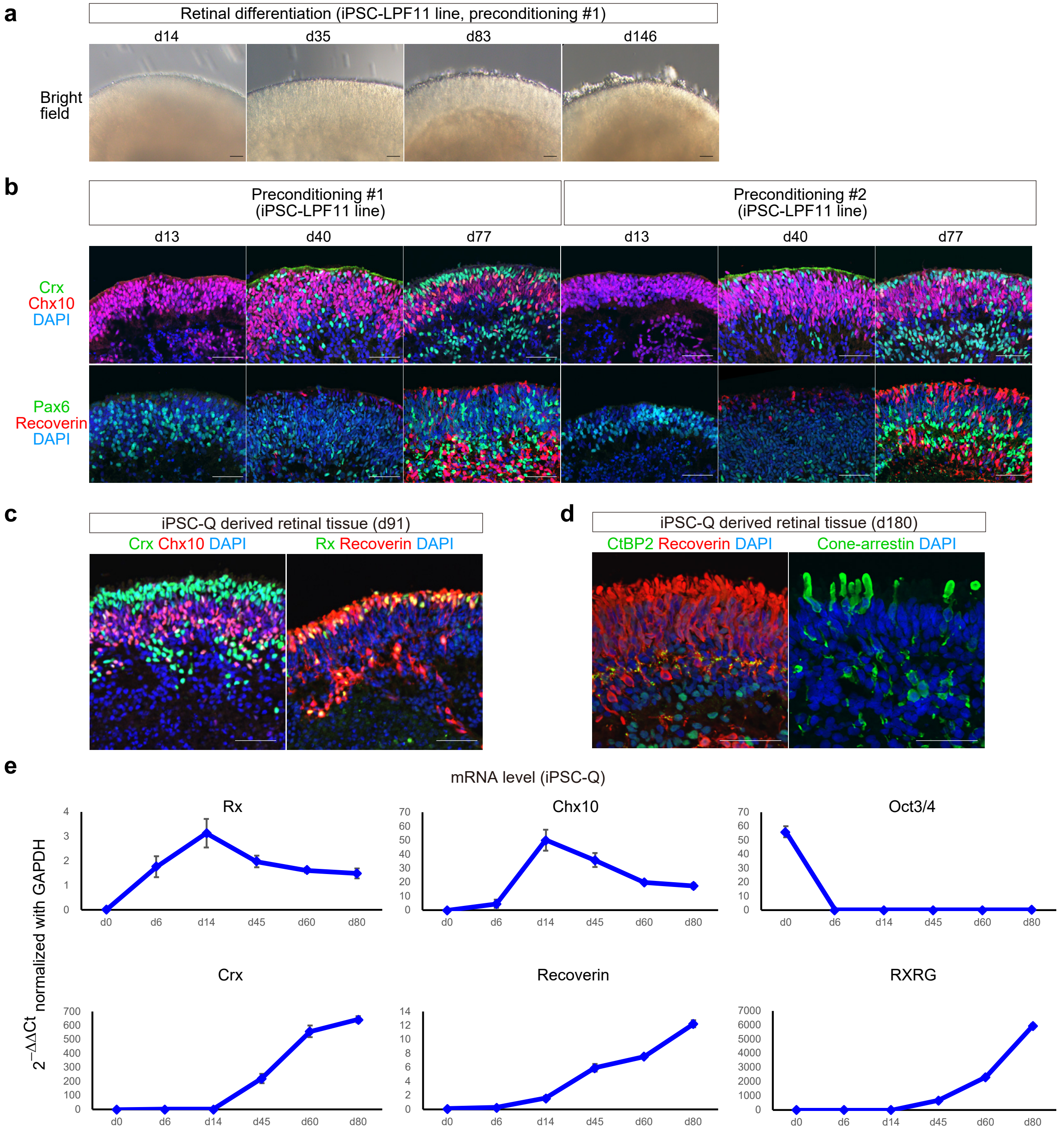


**Supplementary Figure 1. Self-organizing culture of human iPSCs to generate the 3D-retina and dissected retinal sheet (supplementary to Fig. 1).**

**a** Bright-field view of iPSC-LPF11-derived cell aggregate containing retinal tissue on days 14, 35, 83, and 146. Note that the morphologies of the iPSC-LPF11-derived retinal tissue in (**a**) and the iPSC-S17-derived retinal tissue in Fig. 1b were similar at each time point. Scale bar: 100 µm.

**b** Immunostaining of iPSC-LPF11-derived retinal tissue on days 13, 40, and 77 differentiated from preconditioned iPSCs with preconditioning #1 (SB+SAG; left panels) and #2 (SAG-only; right panels). Crx (green) and Chx10 (red) in upper panels. Pax6 (green) and Recoverin (red) in lower panels. Blue: nuclear staining with DAPI. Scale bar: 50 µm. Note that the expressions of retinal marker genes, Crx, Chx10, Pax6, and Recoverin, at each time point were similar in retinal tissues derived from both preconditioning #1 and #2.

**c,d** Immunostaining of iPSC-Q-derived retinal tissue on days 91 (**c**) and 180 (**d**). Crx (green) and Chx10 (red) in (**c**, left). Rx (green) and Recoverin (red) in (**c**, right). CtBP2 (green) and Recoverin (red) in (**d**, left). Cone-arrestin (green) in (**d**, right). Blue: nuclear staining with DAPI. Scale bar: 50 µm.

**e** Gene expression analysis in iPSC-Q-derived cell aggregates containing retinal tissue on days 0, 6, 14, 45, 60, and 80. mRNA levels were determined by qPCR analysis. Relative mRNA expression was determined by the delta-delta Ct method with *GAPDH* as an endogenous control. Data are presented as mean ± SE ($n$ = 48 aggregates per time point).

### Supplementary Figure 2

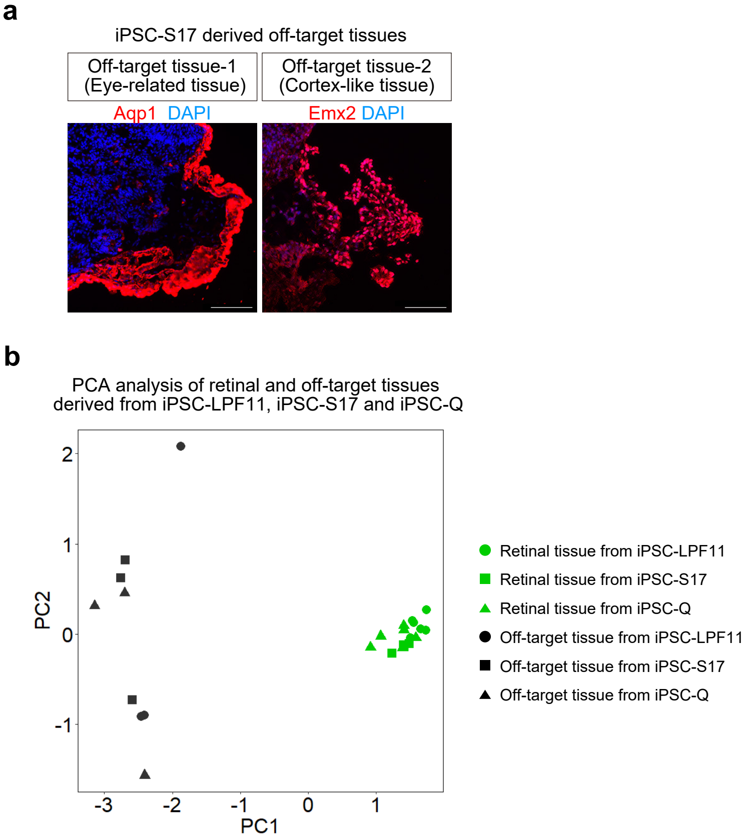


**Supplementary Figure 2. Characterization of major off-target tissues in the retinal differentiation culture (supplementary to Fig. 2).**
**a** Representative IHC images for off-target tissue-1 and -2 derived from iPSC-S17 cells. Immunostaining for Aqp1 (left; red) and Emx2 (right; red) is shown. Blue: nuclear staining with DAPI. Scale bar: 100 µm.
**b** Principal component analysis (PCA) for gene expressions in retinal tissues and off-target tissues derived from iPSC-LPF11, iPSC-S17, and iPSC-Q cells. PC1 and PC2 are shown as scatter plots. The contribution of PC1 and PC2 was 78.2% and 8.8%, respectively.

### Supplementary Figure 3

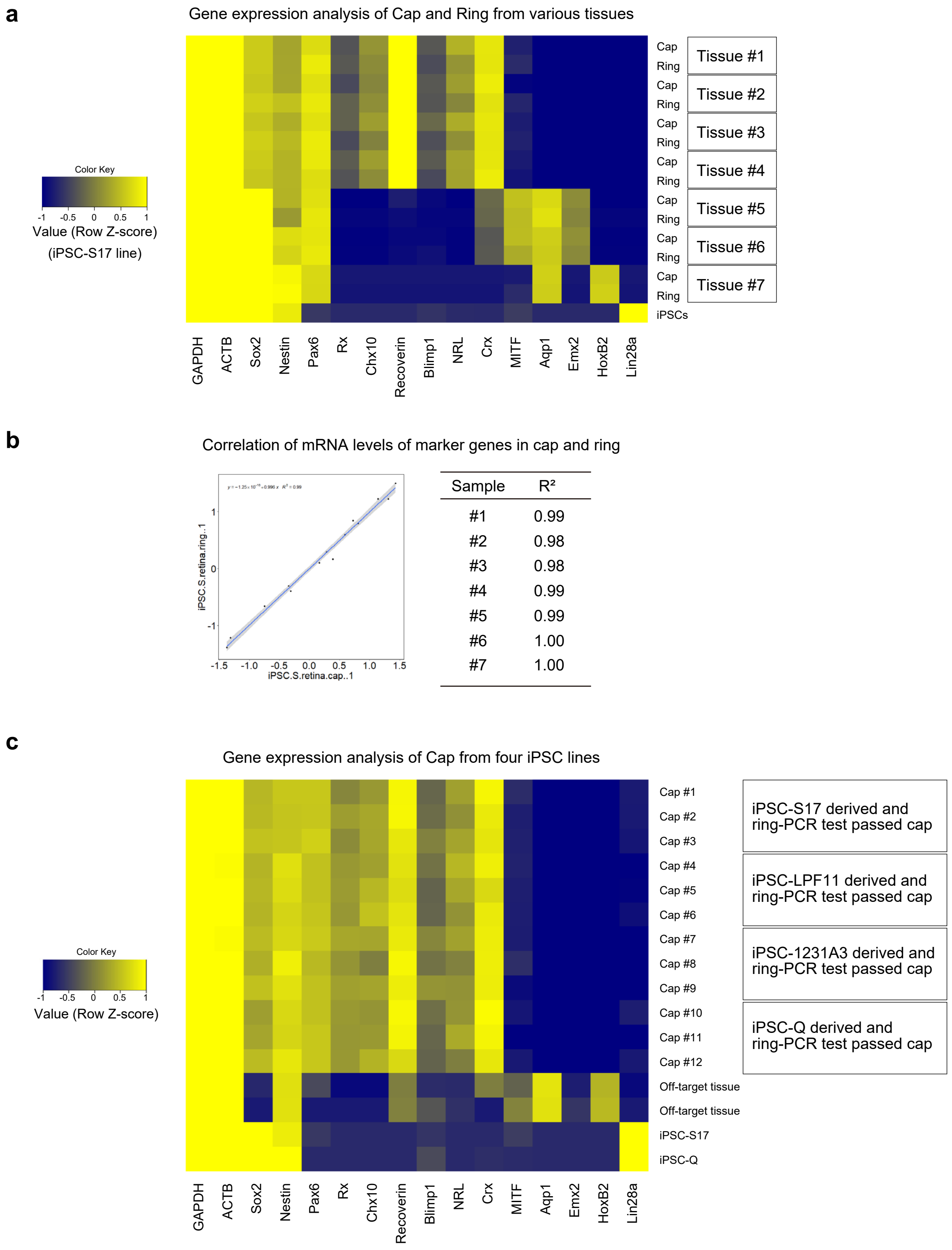


**Supplementary Figure 3. QC method for retinal sheets by testing the gene expression in the surrounding outer tissue-sheet (supplementary to Fig. 3).**
**a** Heat map for gene expressions in caps and rings derived from iPSC-S17 cells. As a control, gene expressions in undifferentiated iPSC-S17 cells are also shown. Gene expressions were measured by qPCR analyses. Row Z-scores were calculated for each cap and ring, and plotted as a heat map.
**b** Representative gene expressions in cap and ring dissected from the same aggregate are shown as a plot with a regression line (left). The coefficient of determination ($R^2$) for the gene expression between each cap and ring is shown (right). #1–#7 correspond to Tissue #1–#7 in (**a**).
**c** Heat map for gene expressions in retinal sheets derived from iPSC-S17, iPSC-LPF11, iPSC-1231A3, and iPSC-Q cells. As a control, gene expressions in off-target tissues derived from iPSC-S17 cells, undifferentiated iPSC-S17 cells, and undifferentiated iPSC-Q cells are also shown.

### Supplementary Figure 4

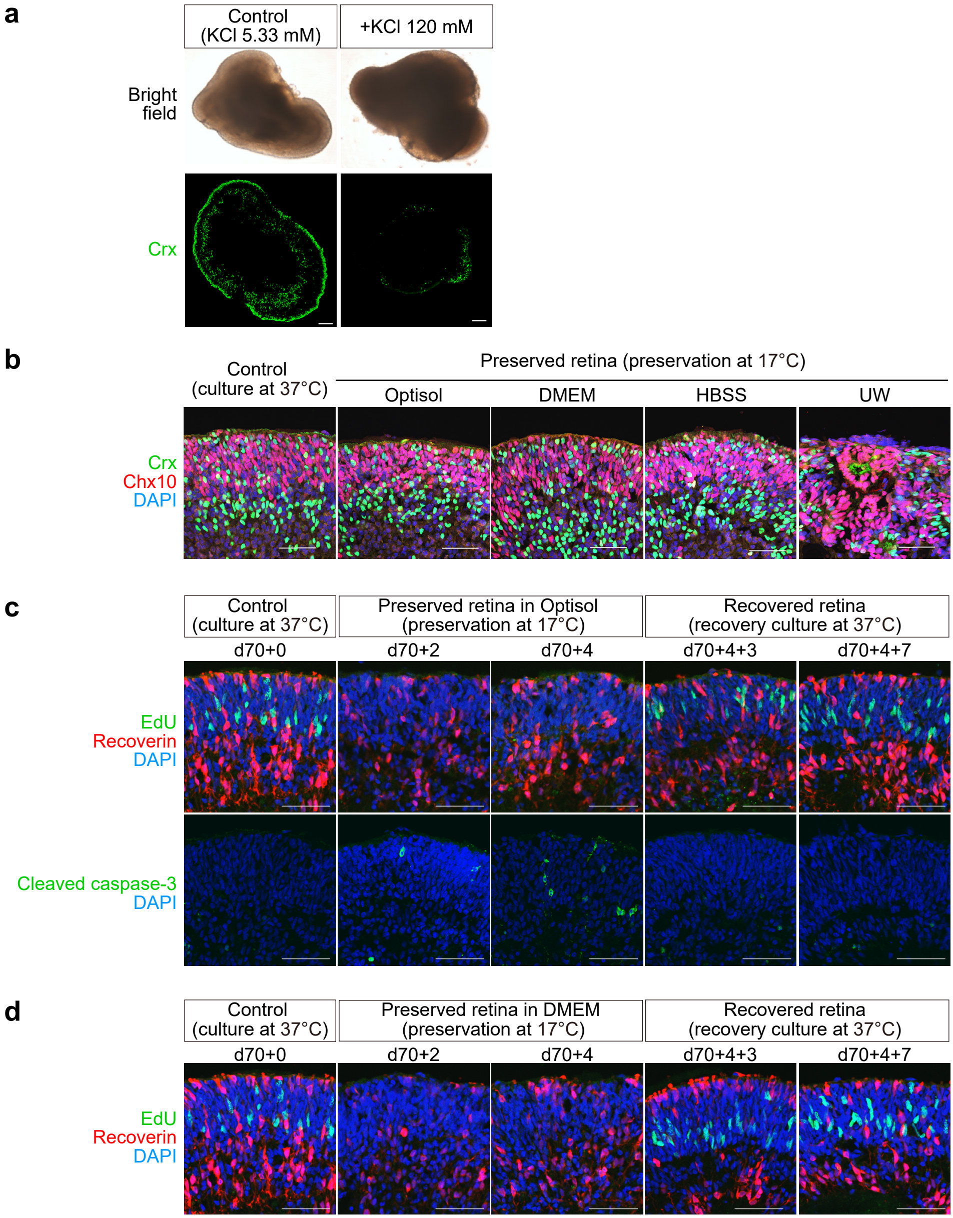


**Supplementary Figure 4. Controlled room temperature non-freezing preservation method (supplementary to Fig. 4).**
**a** Representative bright-field images and immunohistochemical images of retinas preserved in Optisol (Control) and Optisol supplemented with potassium chloride (+KCl 120 mM). After preservation for 3 days at 17 °C, the retinas were cultured for 7 days at 37 °C under 5% $CO_2$ as a recovery culture. Bright-field images are shown in the upper panels and immunohistochemical images for Crx (green) are shown in the lower panels. Scale bar in immunostaining: 100 µm.
**b** Representative immunohistochemical images of retinas preserved in Optisol, DMEM, BSS, and UW. After preservation at 17 °C for 3 days, the retinas were cultured for 7 days at 37 °C under 5% $CO_2$ as a recovery culture. Immunohistochemical images for Crx (green) and Chx10 (red) are shown. Blue: nuclear staining with DAPI. Scale bar: 50 µm.
**c** Representative immunohistochemical images of retinas that were not preserved (d70+0), preserved in Optisol for 2 days (d70+2) and 4 days (d70+4), and preserved in Optisol for 4 days followed by recovery culture for 3 days (d70+4+3) and 7 days (d70+4+7). Immunostaining for EdU (green) and Recoverin (red) are shown in upper panels, and immunostaining for cleaved caspase 3 (green) is shown in the lower panels. Note that the increase in apoptosis or non-apoptotic caspase-3 cleavage during preservation may be transient. DAPI is shown in blue. Scale bar: 50 µm.
**d** Representative immunohistochemical images of retinas that were not preserved (d70+0), preserved in Dulbecco's modified eagle medium (DMEM), basal medium which is extensively used for mammalian cell culture, for 2 days (d70+2) and 4 days (d70+4), and preserved in DMEM for 4 days followed by recovery culture for 3 days (d70+4+3) and 7 days (d70+4+7). Immunostaining for EdU (green) and Recoverin (red) are shown in upper panels. DAPI is shown in blue. Scale bar: 50 µm.

### Supplementary Figure 5

**a**

Group composition

| | 13W | 26W | 52W | 78W |
|---|---|---|---|---|
| Intact | 2 | 0 | 2 | 4 |
| Sham | 0 | 4 | 4 | 4 |
| Transplanted | 4 | 6 | 7 | 4 |

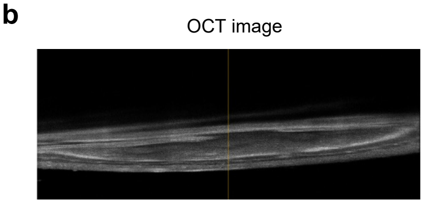


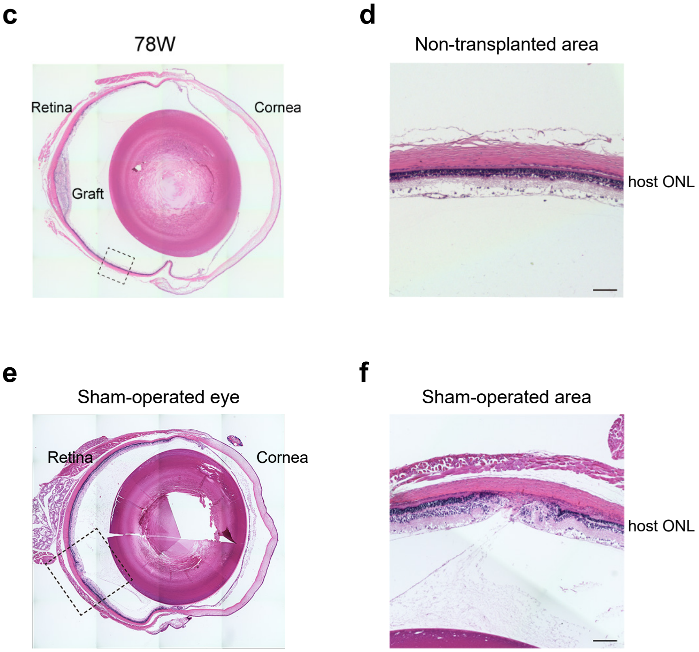


**Supplementary Figure 5. *In vivo* tumorigenicity study by subretinal transplantation in nude rats (supplementary to Fig. 5).**
**a** Group composition of the nude rats in the tumorigenicity study. Intact: control nude rat without surgery (*n* = 8). Sham: control nude rat with subretinal injection of vehicle control (*n* = 12). Transplanted: nude rat with subretinal transplantation of a single iPSC-Q-derived retinal sheet (*n* = 21).
**b** OCT image of a rat retina after subretinal transplantation to confirm the location of the retinal sheet in the subretinal space.
**c, d** HE staining of a transplanted eye section at 78 weeks after transplantation. The magnification image of control non-transplanted area is shown in (**d**). Note that the retina in the control non-transplanted area contained outer nuclear layer.
**e, f** HE staining of a sham-operated eye section at 78 weeks after surgery. The magnification image of sham-operated area is shown in (**f**). Note that the lamination of host outer nuclear layer is affected in the sham-operated area.
ONL, outer nuclear layer. Scale bars in (**d, f**): 100 µm.

### Supplementary Figure 6

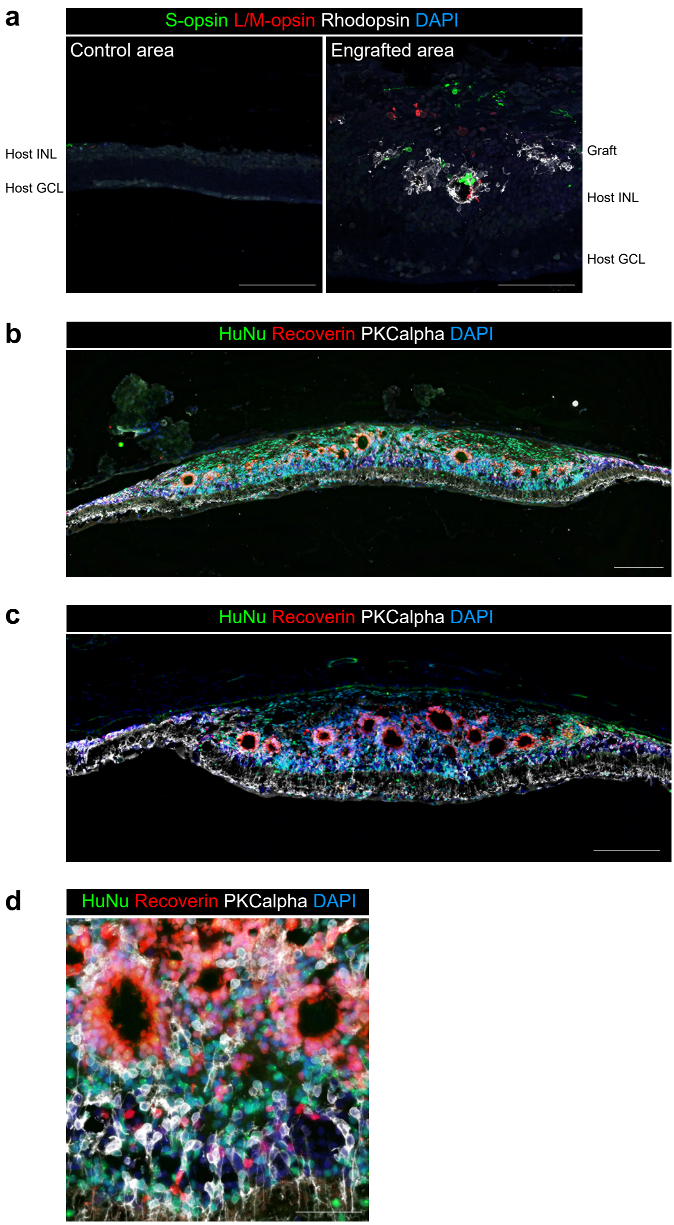


**Supplementary Figure 6. Engraftment and photoreceptor maturation of retinal sheets after subretinal transplantation in RD-nude rats (supplementary to Fig. 6).**

**a** Immunostaining of rat eyes transplanted with iPSC-S17-derived retinal sheets. The magnification images of control non-transplanted area and engrafted area are shown. The retinal sheets were transplanted in the subretinal space of RD-nude rats. The rat retinas were fixed at 263 days after transplantation (341 days after initiation of differentiation). Immunostaining for S-opsin (green), L/M-opsin (red), and Rhodopsin (white) in control non-transplanted area are shown. DAPI is shown in blue.

**b–d** Immunostaining of rat eyes transplanted with iPSC-Q-derived retinal sheets. The retinal sheets were transplanted in the subretinal space of RD-nude rats ($n = 2$). Two rat retinas fixed at 153 days after transplantation (237 days after differentiation) in (**b**) and 217 days after transplantation (301 days after differentiation) in (**c, d**) are shown. Immunostaining for HuNu (green), Recoverin (red) PKCalpha (white) are shown. DAPI is shown in blue.

INL, inner nuclear layer; GCL, ganglion cell layer. Scale bars: 100 µm in (**a**), 200 µm in (**b, c**), 50 µm in (**d**).

### Supplementary Figure 7

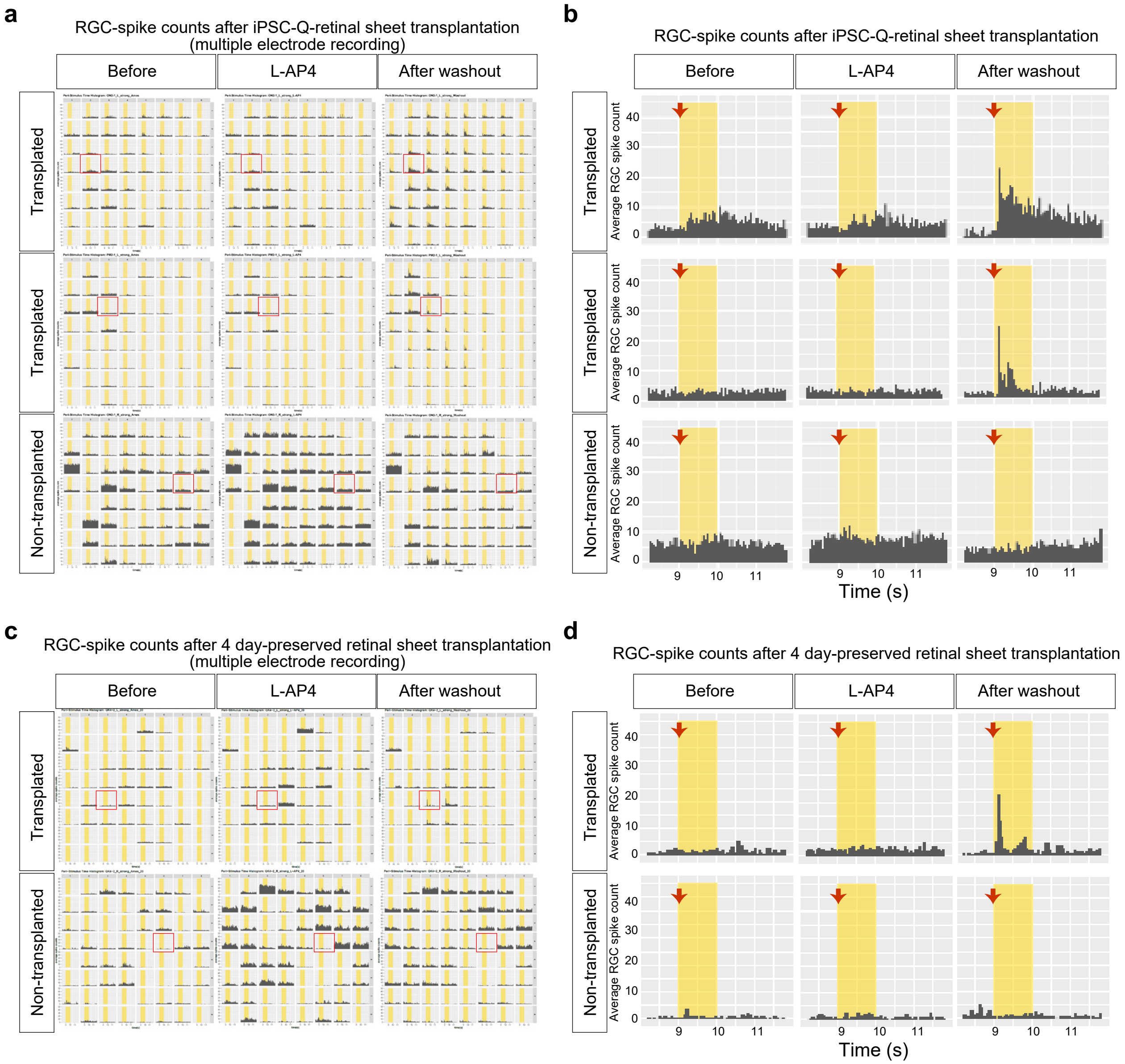


**Supplementary Figure 7. Light responses in transplanted iPSC-retinas by the MEA assay (supplementary to Fig. 7).**
**a–d** *Ex vivo* electrophysiology assay using MEA to evaluate the light responses in transplanted iPSC-retinas. Transplanted and non-transplanted rat retinas were mounted on the MEA electrodes for electrophysiological recording. During the recording, light stimuli were conducted in different intensities (weak, medium, strong), and the rat retinas were incubated in medium (Before), followed by addition of L-AP4 (L-AP4), and washout of L-AP4 (After washout). Peri-stimulus time histograms to examine the RGC-spike counts during strong light stimuli (12.84 log photons/cm²/s) are shown in (**a–d**). The X-axis and Y-axis represent the time and average RGC-spike counts, respectively. Recording data for iPSC-retinal sheet-transplanted rat retinas (Transplanted) and non-transplanted rat retinas (Non-transplanted) in the three conditions (Before, L-AP4, After washout) are shown. Multiple electrode recording data are shown in (**a, c**) and representative data (red box in **a, c**) are shown in (**b, d**). Yellow box in (**a–d**): light stimuli. Red arrows in (**b, d**): start timing of light stimuli.
**a, b** Two results obtained for RD-nude rat retinas transplanted with an iPSC-Q-derived retinal sheet after RT-preservation for 3 days in Optisol.
**c, d** One result obtained for an RD-nude rat retina transplanted with an iPSC-S17-derived retinal sheet after RT-preservation for 4 days in medium.

### Supplementary Figure 8

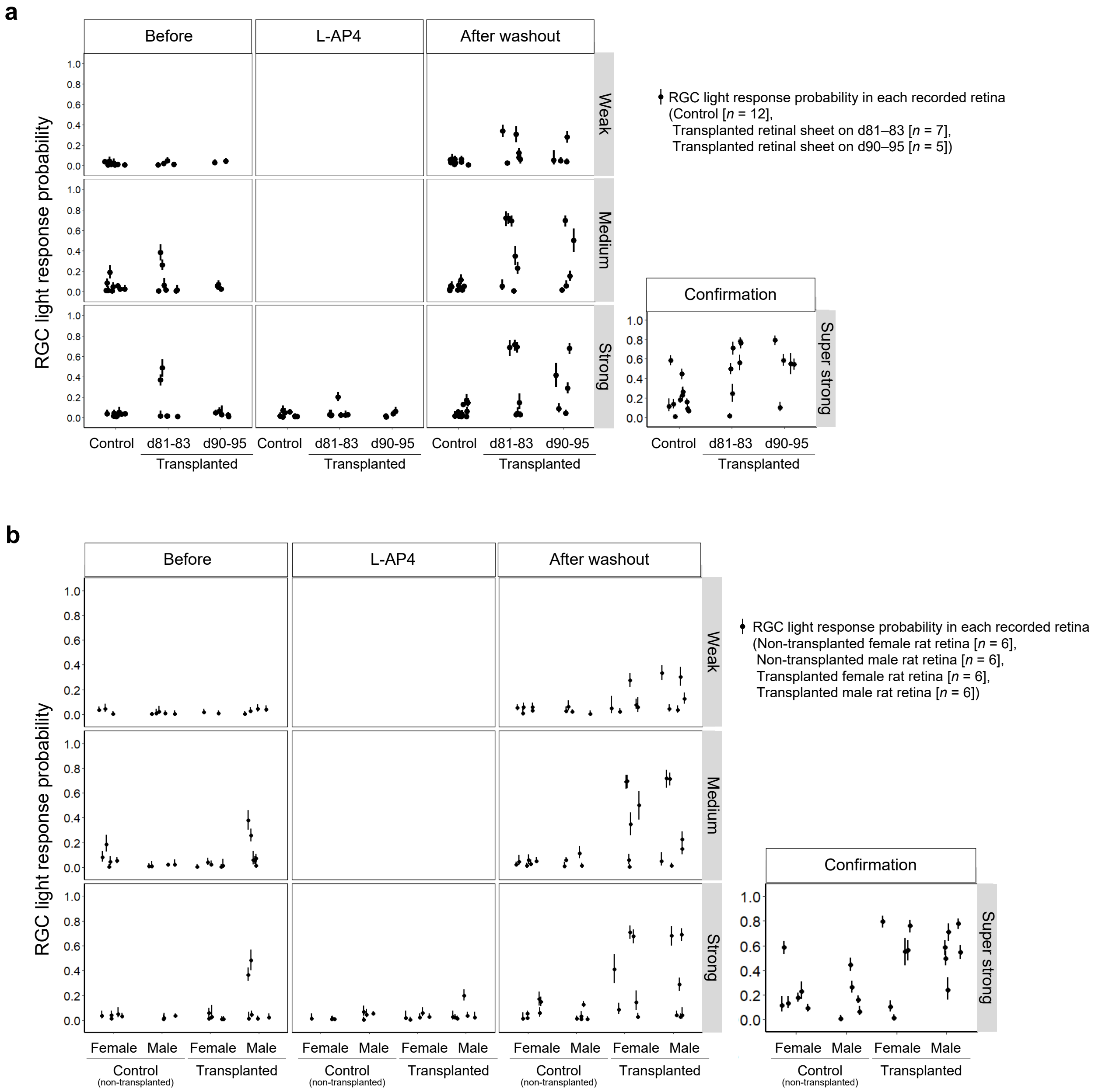


**Supplementary Figure 8. RGC light response probabilities in transplanted iPSC-retinas by the MEA assay (supplementary to Fig. 7).**
**a, b** *Ex vivo* electrophysiology assay using MEA to evaluate the light responses in transplanted iPSC-retinas. Rat retinas were subjected to MEA analysis to evaluate the RGC light response probability after stimulation. RGC light responses in non-transplanted rat retinas (Control; *n* = 12), rat retinas transplanted with an iPSC-S17-derived retinal sheets cultured for days 81–83 (Retinal sheet on d81–83; *n* = 7), and rat retinas transplanted with an iPSC-S17-derived retinal sheets cultured for days 90–95 (Retinal sheet on d90–95; *n* = 5) are shown in (**a**). RGC light responses in female (*n* = 6) and male (*n* = 6) non-transplanted rat retinas (Control) and female (*n* = 6) and male (*n* = 6) rat retinas transplanted with an iPSC-S17-derived retinal sheets (Transplanted) are shown in (**b**). Dots and bars show the per sample (recorded rat retina) summary of the collected data. Dot: RGC light response probability estimated by statistical modeling (Bayesian statistical inference with Markov Chain Monte Carlo sampling). Bar: 89% confidence interval.

### Supplementary Figure 9

| hPSC line | hESC-KhES1 | iPSC-1231A3 | iPSC-LPF11 | iPSC-S17 | iPSC-QHJI01s04 |
|---|---|---|---|---|---|
| Self-organization culture | Refs. 1,2,3 | Ref. 3 | Ref. 3, Fig. 2b, S1a,b | Fig. 1a-c, 2b | Fig. S1c,d |
| Quality | Ref. 7 | Fig. 2e, S3c* | Fig. 2a-c, S3c* | Fig. S3a, S3c* | Fig. S3c* |
| Engraftment and maturation | Refs. 4,5 | Refs. 3,6 | n.d. | Fig. 6* | Fig. S6* |
| Preclinical in vivo function (*ex vivo* MEA assay) | Refs. 5,7 | Refs. 6 | n.d. | Fig. 7*, S7*, S8* | Fig. S7* |
| *In vivo* tumorigenicity | n.d. | n.d. | n.d. | n.d. | Fig. 5*, S5* |

**Supplementary Figure 9. Five human PSC lines and their preclinical data in the previous and present studies.**
Figures in the present study and the references of previous reports are shown to summarize which experimental data have been obtained from five human PSC lines, hESC-KhES1, iPSC-1231A3, iPSC-LPF11, iPSC-S17, and iPSC-QHJI01s04.

In previous studies, we have developed self-organizing culture to generate 3D-retina from hESC-KhES1, iPSC-1231A3 cells, and iPSC-LPF11 lines (1, 2, 3). We have previously shown that the retinal sheets derived from these three ESC and iPSC lines could become engrafted in retinal degenerative model mice (5), rats (3, 6), and non-human primates (4, 6). In the present study, we at first improved self-organizing culture to generate 3D-retina from new iPSC lines, iPSC-S17 and iPSC-Q (Fig. 1 and Supplementary Fig. 1). We confirmed that both iPSC-S17 and iPSC-Q self-formed multilayered retinal tissue in the retinal differentiation culture as previously observed in retinal differentiation from iPSC-1231A3 and iPSC-LPF11. This indicated that our self-organizing culture was a robust method to generate 3D-retinas from various PSC lines.

Then, we established the QC method including the ring-PCR test. We investigated the gene expression of the retinal sheets, which passed ring-PCR test, and found that retinal sheets derived from iPSC-LPF11, iPSC-1231A3, iPSC-S17, and iPSC-Q showed similar retinal gene expression patterns (Supplementary Fig. 3c). This demonstrated that the combination of self-organizing culture and ring-PCR test is the robust method to generate the retinal sheet of controlled quality from various PSC lines.

We also have developed RT-preservation method. We confirmed that preserved retinal sheets derived from iPSC-S17 and iPSC-Q become engrafted in RD-nude rats and differentiated into mature photoreceptors (Fig. 6), as observed in our previous studies using hESC-KhES1 and iPSC-1231A3 (3, 6, 7). Importantly, the preserved retinal sheets derived from both iPSC-S17 and iPSC-Q had a potential to improve light responsiveness evaluated by ex vivo electrophysiology assay (Fig. 7 and Supplementary Figs. 7, 8), as observed in our previous studies (6, 7).

Collectively, these results demonstrated that the self-organizing culture, ring-PCR test, and RT-preservation method can be applicable to various PSC lines to generate the retinal sheets with controlled quality.


n.d.: not determined.
*Retinal sheet obtained by ring-PCR and RT-preservation methods.
